# Polygenic viral factors enable efficient mosquito-borne transmission of African Zika virus

**DOI:** 10.1101/2025.01.23.634482

**Authors:** Shiho Torii, Jennifer S. Lord, Morgane Lavina, Matthieu Prot, Alicia Lecuyer, Cheikh T. Diagne, Oumar Faye, Ousmane Faye, Amadou A. Sall, Michael B. Bonsall, Etienne Simon-Lorière, Xavier Montagutelli, Louis Lambrechts

## Abstract

Zika virus (ZIKV) is a mosquito-borne flavivirus primarily transmitted among humans by *Aedes aegypti*. Over the past two decades, it has caused significant outbreaks associated with birth defects and neurological disorders. Phylogenetically, ZIKV consists of two main genotypes referred to as the African and Asian lineages, each exhibiting distinct biological properties. African lineage strains are transmitted more efficiently by mosquitoes, but pinpointing the genetic basis of this difference has remained challenging. Here, we address this question by comparing recent African and Asian strains using chimeric viruses, in which segments of the parental genomes are swapped. Our results show that the structural genes from the African strain enhance viral internalization, while the non-structural genes improve genome replication and infectious particle production in mosquito cells. *In vivo* mosquito transmission is most significantly influenced by the structural genes, although no single viral gene alone determines this effect. Additionally, we develop a stochastic model of *in vivo* viral dynamics in mosquitoes that mirrors the observed patterns, suggesting that the primary difference between the African and Asian strains lies in their ability to traverse the mosquito salivary glands. Overall, our findings suggest that the polygenic nature of ZIKV transmissibility has prevented Asian lineage strains from achieving the same epidemic potential as African lineage strains, underscoring the importance of lineage-specific adaptive landscapes in shaping ZIKV evolution and emergence.

## Introduction

Zika virus (ZIKV) is an emerging mosquito-borne orthoflavivirus of significant global public health concern. It is primarily transmitted among humans by *Aedes aegypti* mosquitoes, which are found in tropical and sub-tropical regions worldwide (*1, 2*). ZIKV gained widespread international attention during the major 2015-2016 outbreak, which affected over 1.6 million people across more than 70 countries in the Pacific and the Americas (*3–5*). While most human infections are either asymptomatic or result in mild symptoms, ZIKV can lead to severe neurological disorders, such as Guillain-Barré syndrome, and congenital abnormalities, including microcephaly, in fetuses of infected pregnant women (*6–10*). Given the absence of specific antiviral treatments or vaccines, understanding ZIKV transmission dynamics is critical for developing effective prevention and control strategies.

First identified in 1947 in a sentinel rhesus monkey from the Zika Forest in Uganda (*11*), ZIKV circulated largely undetected for the next six decades, likely within sylvatic cycles between forest-dwelling mosquitoes and non-human primates, with only 14 sporadic reports of human infections in parts of Africa and Asia (*6, 12, 13*). The first documented outbreak occurred in 2007 on the Pacific Island of Yap, Micronesia (*13*), followed by rapid spread to other Pacific islands and eventually the Americas. Phylogenetically, ZIKV is divided into two main genotypes sharing ∼90% nucleotide identity, referred to as the African and Asian lineages, which diverged in the early 19^th^ century (*9, 14–16*). All recorded ZIKV outbreaks, including a recent one in Cape Verde, have been caused by Asian lineage strains, whereas African lineage strains have only been detected in 5 countries in Africa (*13, 17, 18*). Furthermore, accumulating evidence suggests that African ZIKV strains, although not implicated in human outbreaks to date, exhibit higher infectivity and pathogenicity compared to Asian strains in various *in vitro* and *in vivo* human models (*16, 19–27*).

The dramatic global emergence of ZIKV in the Pacific and the Americas spurred extensive research into the genetic mutations that may have facilitated its spread in the human population, such as mutations enhancing human tropism, pathogenesis, or transmission by *Ae. aegypti* (*28*). The ZIKV genome consists of a positive-sense, single-stranded RNA that encodes a single open reading frame, flanked by 5’ and 3’ untranslated regions (UTRs). This open reading frame translates into a polyprotein, which is post-translationally cleaved into three structural proteins (SPs) – capsid (C), precursor membrane (prM), and envelope protein (E) – and eight non-structural proteins (nSPs) – NS1, NS2A, NS2B, NS3, NS4A, 2K, NS4B, and NS5 (*29*). Reverse genetics studies comparing the two ZIKV lineages showed that introducing Asian SPs into an African strain reduces viral infectivity in human epithelial and neuronal cells (*30*), while introducing African SPs into an Asian strain increases pathogenicity in mice (*31*). Additionally, African strains demonstrate greater transmissibility in *Ae. aegypti* mosquitoes (*27, 32–35*), yet the specific viral genetic factors contributing to this difference remain unclear. Functional studies have shown that two point mutations that arose during ZIKV emergence since 2013 – A188V in NS1 and T106A in C – enhance the replication of Asian strains in *Ae. aegypti* mosquitoes (*36, 37*). Intriguingly, the same transmission-adaptive variants are also present in sylvatic African strains, suggesting they could be ancestral to the virus (*38*). However, collectively these mutations do not account for the full difference in mosquito transmissibility observed between the Asian and African lineages (*38*), implying that other genetic factors have constrained Asian lineage strains from achieving the same epidemic potential as African lineage strains.

Here, we conducted a comprehensive comparison of recent African and Asian ZIKV strains to systematically identify the viral genetic factors that influence the efficiency of mosquito-borne transmission. Using chimeric viruses, we assessed the contribution of each region of the viral genome to differences in viral propagation in mosquito cells *in vitro* and mosquito transmissibility *in vivo*. Finally, we obtained mechanistic insights into ZIKV transmissibility by integrating experimental data with a stochastic model of *in vivo* viral dynamics.

## Results

### Engineered ZIKV strains recapitulate the phenotypes of natural isolates in mosquitoes

To understand the viral genetic factors impacting mosquito transmissibility, we selected, because of their distinct levels in transmission efficiency (*27*), recent ZIKV strains from Senegal and Thailand, representing the African and Asian lineages, respectively. To minimize the impact of natural genetic variability, we used reverse genetics to transform the original virus isolates, iSenegal and iThailand, into engineered strains, which we designated as rSenegal and rThailand, respectively (Figure S1A). To assess whether the strains generated by reverse genetics recapitulated the phenotypes of their natural counterparts, we exposed *Ae. aegypti* mosquitoes from Colombia to each of these viruses via artificial infectious blood meals (Figure S1B). At 7 and 14 days post blood meal, we measured the prevalence of infection and systemic viral dissemination by RT-PCR and evaluated ZIKV transmission potential by detecting the presence of infectious virus in salivary secretions (Figure S2, Table S3). Both rSenegal and rThailand exhibited a similar prevalence of infection, systemic dissemination, and viral presence in saliva compared to their natural counterparts. In line with our earlier study (*27*), both iSenegal and rSenegal demonstrated significantly higher dissemination prevalence and greater transmission efficiency than iThailand and rThailand across the tested time periods (Figure S2B-C). These findings confirm the comparability of the ZIKV strains generated by reverse genetics to the natural isolates in terms of their infection dynamics in mosquitoes. Consequently, we used rSenegal and rThailand as parental strains for all subsequent experiments (Figure S1B).

### Superior growth kinetics of the rSenegal strain reflect higher efficiency of viral internalization and genome replication in mosquito cells

We first examined the growth kinetics of rSenegal and rThailand strains in mosquito (C6/36) cells *in vitro*. Over the four-day time course, the rSenegal strain consistently showed higher titers compared to rThailand (Figure 1A). To understand why these strains produced different levels of infectious particles, we performed several functional assays. We evaluated viral attachment by measuring membrane-bound viral RNA levels after one-hour viral attachment on ice. This experiment revealed that the rThailand strain had higher attachment efficiency than the rSenegal strain (Figure 1B), thus failing to explain the superior growth kinetics of the rSenegal strain. We also analyzed viral internalization efficiency by measuring relative intracellular viral RNA at 3 hours post infection (h.p.i.) normalized to the attached virus particles (Figure 1C). The validity of the assay was verified with Dynasore, an inhibitor of clathrin-mediated endocytosis, which is considered the primary internalization pathway of orthoflaviviruses (*39–42*). Internalization efficiency was lower under Dynasore treatment (Figure 1C, left panel) and was significantly higher for the rSenegal strain than for the rThailand strain (Figure 1C, right panel), suggesting that differences in internalization efficiency contribute to the higher infectious particle production of the rSenegal strain. Next, we monitored the replication of ZIKV genomic and antigenomic RNA over 24 hours using strand-specific RT-qPCR (Figure 1D). Both strains showed detectable levels of ZIKV antigenomic RNA from 12 h.p.i., with higher replication efficiency in the rSenegal-infected cells detected at 18 h.p.i. (Figure 1D, left panel). The relative viral genomic RNA level was also significantly higher in rSenegal-infected cells from 15 to 24 h.p.i. (Figure 1D, right panel). This result indicates that a higher efficiency of viral genome replication by the rSenegal strain may also contribute to the higher production of infectious virus particles. We also explored the ratio of infectious virus titers to viral RNA copies to assess the release of non-infectious immature or defective virus particles (*43–46*) (Figure 1E). Although this ratio increased over time for both strains, it was significantly lower in rSenegal-infected cells at 48 h.p.i., suggesting a slightly higher proportion of immature or defective virus particles produced by the rSenegal strain. By monitoring the decay of infectious virus titers over time at 28°C, we found that there was no detectable difference in the stability of infectious virus particles between the rSenegal and rThailand strains (Figure 1F). Finally, we observed that the rSenegal strain significantly reduced cellular ATP levels compared to the rThailand strain at 96 h.p.i. (Figure 1G). Together, these results indicate that the more efficient virus particle production of the rSenegal strain in mosquito cells reflects enhanced virus internalization and viral genome replication efficiency, despite increased cell toxicity.

**Figure 1.**
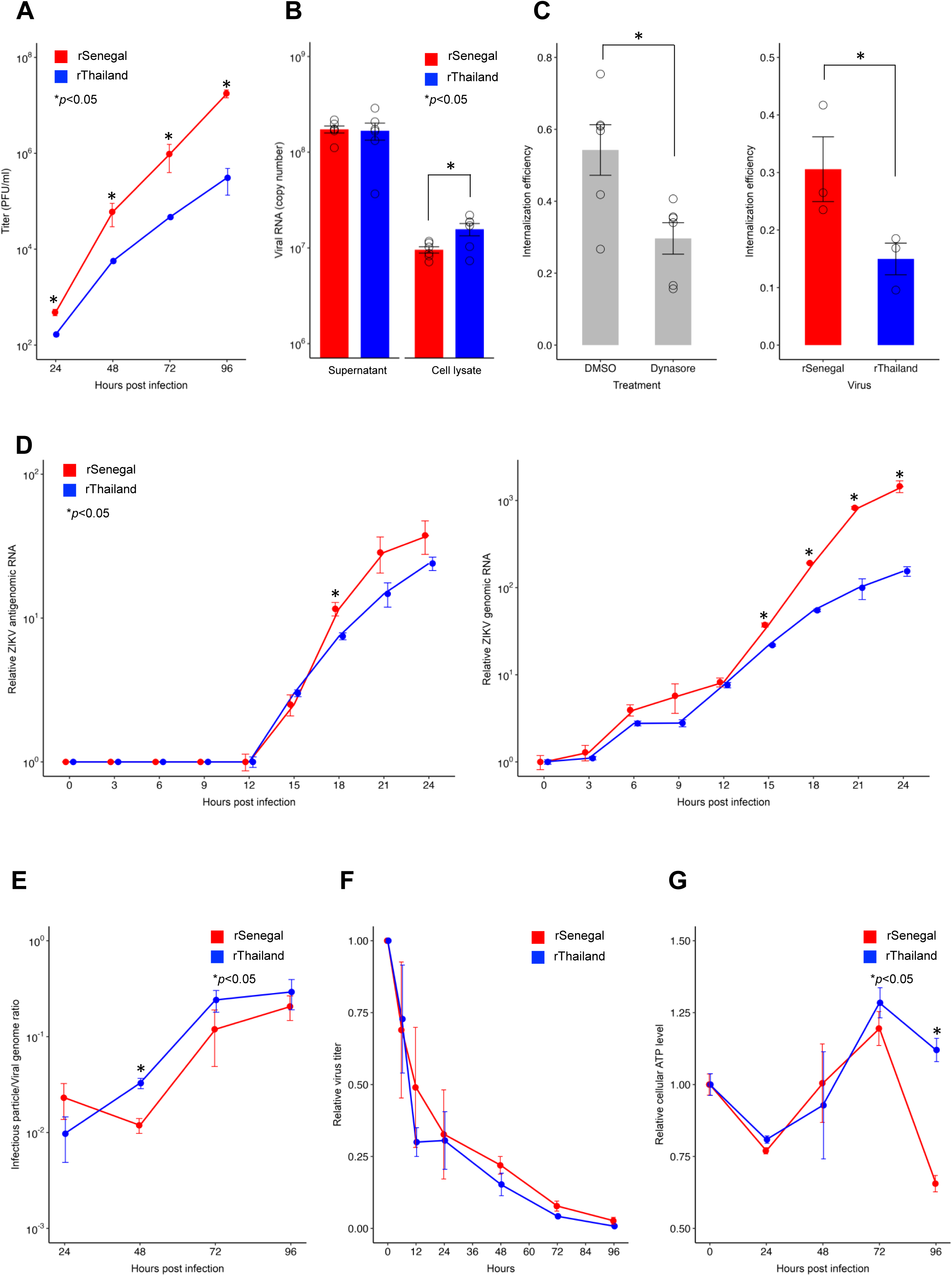
Superior growth kinetics of the rSenegal strain are associated with enhanced efficiency of viral internalization and genome replication in mosquito cells. (**A**) Viral growth kinetics of the rSenegal and rThailand strains were assessed by determining infectious virus titers in the supernatant of ZIKV-infected C6/36 cells (MOI of 0.01) by plaque assay. (**B**-**D**) The efficiency of viral attachment (**B**), internalization (**C**), and genome replication (**D**) was measured in C6/36 cells infected at a MOI of 1. (**B**) Viral RNA levels bound to the membrane and in the supernatant were quantified one hour post attachment by incubating cells on ice. (**C**) Internalization efficiency was analyzed by quantifying viral RNA relative to *Actin* mRNA at 3 hours post infection (h.p.i.) following protease E treatment. Internalization was normalized to attached viral RNA levels, in cells infected with the rSenegal strain treated with DMSO or Dynasore (left panel) and in cells infected with the rSenegal or the rThailand strains (right panel). (**D**) Genome replication was assessed over 24 hours by normalizing ZIKV antigenomic RNA levels to those at 12 h.p.i. (left panel) and genomic RNA levels to those at 0 h.p.i. (right panel). (**E**) The virus titer-to-viral genome ratio was analyzed over time by infecting C6/36 cells at an MOI of 0.01 and measuring infectious titers by plaque assay and viral genome copy number by RT-qPCR in supernatants. (**F**) Viral decay rate was determined by monitoring infectious titer over time at 28°C, normalized to titer at 0 hours. (**G**) The cellular ATP levels of ZIKV-infected C6/36 cells was assessed over time using the CellTiter-Glo assay and normalized to the levels at 0 h.p.i. (**A**-**G**) Data are presented as mean ±SEM from 3-6 biological replicates. Statistical significance was determined using Student’s *t*-test (\**p* <0.05; ns: non-significant). Lines and bars are color-coded according to the virus strain.

### Structural genes enhance viral internalization, and non-structural genes increase genome replication of rSenegal strain relative to rThailand strain in mosquito cells

To investigate the genetic determinants responsible for differences in mosquito transmissibility between the rSenegal and rThailand strains, we analyzed the full-length genome sequences of these strains (Figure S1C). Since comparative sequence analysis identified 102 amino-acid differences and 1216 nucleotide differences without specific clustering, we constructed a first set of chimeric viruses by swapping SPs, nSPs, and UTRs between the parental strains by reverse genetics (Figure 2A). We observed significant differences in plaque size on a mammalian (Vero E6) cell monolayer between the chimeric viruses. Adding the rSenegal SPs into the rThailand backbone increased plaque size, whereas adding the rThailand SPs into the rSenegal backbone decreased it (Figure 2B). The rThailand nSPs introduced into the rSenegal backbone led to smaller plaques but there was no detectable effect of the reciprocal replacement. In contrast, introducing rSenegal UTRs into the rThailand backbone reduced plaque size, while doing so in the opposite direction showed no significant change (Figure 2B). These results suggest that SPs, nSPs, and UTRs, influence plaque size, although their effects vary depending on the backbone strain. We then assessed the growth kinetics of the chimeric viruses in mosquito cells. Chimeric viruses using the rSenegal strain as a backbone showed that introducing either SPs or nSPs from the rThailand strain significantly reduced virus titers compared to the parental rSenegal strain (Figure 2C, left panel). For chimeric viruses using the rThailand strain as a backbone, replacement of the nSPs increased virus titers, particularly from 24 to 72 h.p.i. (Figure 2C, right panel). These results indicate that swapping either SPs or nSPs influence growth kinetics but with a different magnitude depending on the backbone strain, whereas UTRs do not contribute significantly.

**Figure 2.**
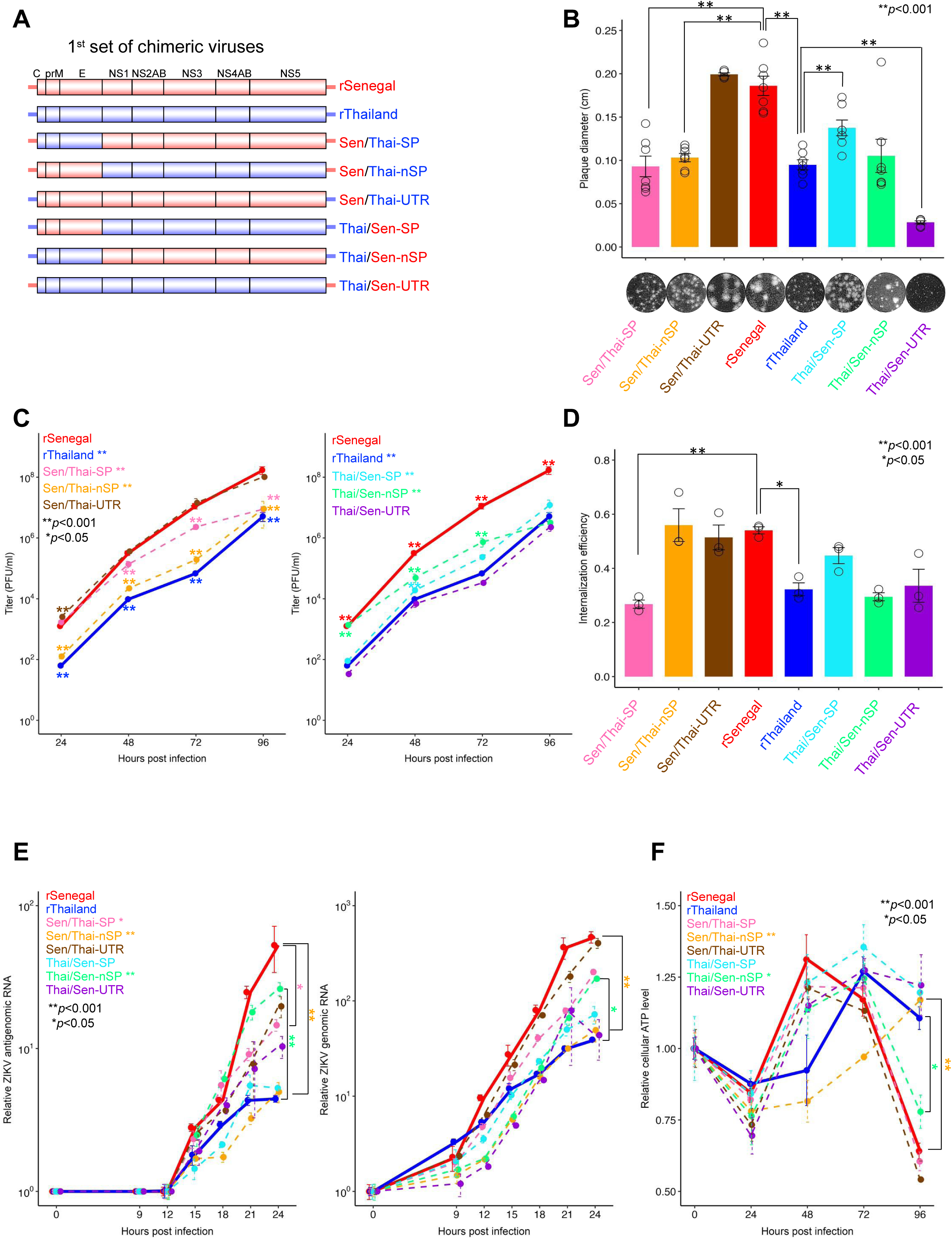
Structural genes enhance viral internalization, and non-structural genes improve genome replication of rSenegal strain relative to rThailand strain in mosquito cells. (**A**) Schematic representation of the first set of chimeric viruses. (**B**) Plaque morphologies of chimeric viruses on Vero E6 cell monolayers. Plaque diameters were measured using ImageJ software for quantification. (**C**) Viral growth kinetics of chimeric viruses with rSenegal (left panel) or rThailand (right panel) backbones were determined by measuring infectious titers from supernatants of ZIKV-infected C6/36 cells (MOI = 0.01) by plaque assay. (**D**-**E**) The efficiency of viral internalization (**D**) and genome replication (**E**) was analyzed in C6/36 cells infected with ZIKV at an MOI of 1. Viral RNA levels were assessed by calculating the ratio of viral RNA to *Actin* expression at 3 h.p.i. for internalization (**D**) and over 24 hours for genome replication (**E**) following protease E treatment. Internalization efficiency (**D**) was determined by normalizing viral RNA at 3 h.p.i. to levels of initially attached viruses. Genome replication (**E**) was evaluated by normalizing ZIKV antigenomic RNA levels at each time point to 12 h.p.i. (left panel) and genomic RNA levels to baseline levels at 0 h.p.i. (right panel). (**F**) The cellular ATP levels following ZIKV infection was assessed over time using the CellTiter-Glo assay and normalized to the levels at 0 h.p.i. (**B**-**F**) Data are presented as mean ±SEM from 3-7 biological replicates. Statistical analysis was performed using one-way ANOVA with Dunnett’s test (\**p* <0.05; \*\**p* <0.01; ns: non-significant). Lines and bars are color-coded to represent the different chimeric viruses, and the parental strains are depicted with thicker lines.

To identify the genomic regions underlying the distinct growth kinetics of the parental strains in mosquito cells, we examined the internalization efficiency of chimeric viruses. The only replacement that significantly influenced internalization efficiency relative to the parental strain was substituting rThailand SPs in the rSenegal strain (Figure 2D). We next compared the replication efficiency of chimeric viruses by measuring the production of ZIKV antigenomic and genomic RNAs. Swapping nSPs between parental strains resulted in significant differences in the production of both viral antigenomic RNA (Figure 2E, left panel) and genomic RNA (Figure 2E, right panel), indicating that nSPs play a crucial role in viral genome replication. Replacing the SPs from the rThailand strain also decreased the production of viral antigenomic RNA (Figure 2E, left panel). Finally, we found significant changes in cellular ATP levels following replacement of nSPs in both directions at 96 h.p.i. (Figure 2F). Taken together, these findings show that differences in growth kinetics between the rSenegal and rThailand strains primarily reflect the effect of SPs on viral internalization and the effect on nSPs on viral genome replication. The nSPs from the rSenegal strain also lead to higher cell toxicity than the nSPs from the rThailand strain.

### Both structural and non-structural genes collectively underlie differences in mosquito transmission between rSenegal and rThailand strains

To investigate the viral genetic basis of transmissibility in mosquitoes *in vivo*, we first assessed the infectivity of the chimeric viruses when delivered through an infectious blood meal. We orally exposed *Ae. aegypti* mosquitoes from Colombia to differing virus doses as outlined in Table S3 and estimated the 50% oral infectious dose (OID_50_) for each chimeric virus from the dose-response curves (Figure S3). The rSenegal strain had a significantly lower OID_50_ estimate than the rThailand strain and introducing either the SPs or the nSPs from the rThailand strain into the rSenegal strain increased the OID_50_ estimates, whereas the rThailand UTRs did not have a detectable impact (Figure S3B). Conversely, introducing the SPs, nSPs, or UTRs from the rSenegal strain into the rThailand strain did not significantly change the OID_50_ estimates. These findings suggest that both SPs and nSPs are required to confer the rSenegal strain a higher infectivity in mosquitoes relative to the rThailand strain. They also show that most chimeric viruses achieve 80-100% infection prevalence when the blood meal titer is >10^6^ PFU/ml. Given this, we chose 2×10^6^ PFU/ml as the standard infectious dose for subsequent experiments.

To assess the mosquito transmissibility of the chimeric viruses, we exposed *Ae. aegypti* mosquitoes from Colombia to each of the viruses via artificial infectious blood meals containing 2×10^6^ PFU/ml. Actual blood meal titers varied from 1.3 to 4.0×10^6^ PFU/ml, and this variation was factored into our statistical analysis (Table S4). At 7, 10, and 14 days post blood meal, we detected midgut infection and systemic viral dissemination by RT-PCR and evaluated transmission potential by detecting the presence of infectious ZIKV in salivary secretions (Figure S1B). At least 20 mosquitoes per time point were tested for each virus (Table S3). We define infection prevalence as the proportion of blood-fed mosquitoes with a virus-positive body, dissemination prevalence as the proportion of infected mosquitoes with a virus-positive head, and transmission prevalence as the proportion of mosquitoes with a virus-positive head releasing infectious virus in their saliva (Figure S1B). Overall transmissibility is encapsulated in transmission efficiency, which is defined as the proportion of blood-fed mosquitoes with infectious saliva. As expected from the infectivity experiment (Figure S3A), the infection prevalence was 80-100% across viruses and time points (Figure 3A). Dissemination prevalence was also 80-100% across viruses and time points (Figure 3B). Both infection prevalence and dissemination prevalence were significantly influenced by the virus (Table S4), however the magnitude of these differences was modest, ranging on average from 83.3% to 99.0% and from 84.5% to 96.8%, respectively (Figure 3A-B). In contrast, transmission prevalence significantly increased over time and ranged from 0% to ∼50% across viruses and time points (Figure 3C). Notably, the rSenegal strain resulted in significantly higher transmission prevalence than the rThailand strain. Introducing either the SPs or the nSPs from the rThailand strain into the rSenegal strain decreased transmission prevalence, whereas the rThailand UTRs did not have a detectable impact. Conversely, only the SPs from the rSenegal strain increased transmission prevalence when introduced into the rThailand strain. These findings indicate that differences in transmission prevalence between the parental strains are primarily determined by the SPs and, to a lesser extent, the nSPs.

**Figure 3.**
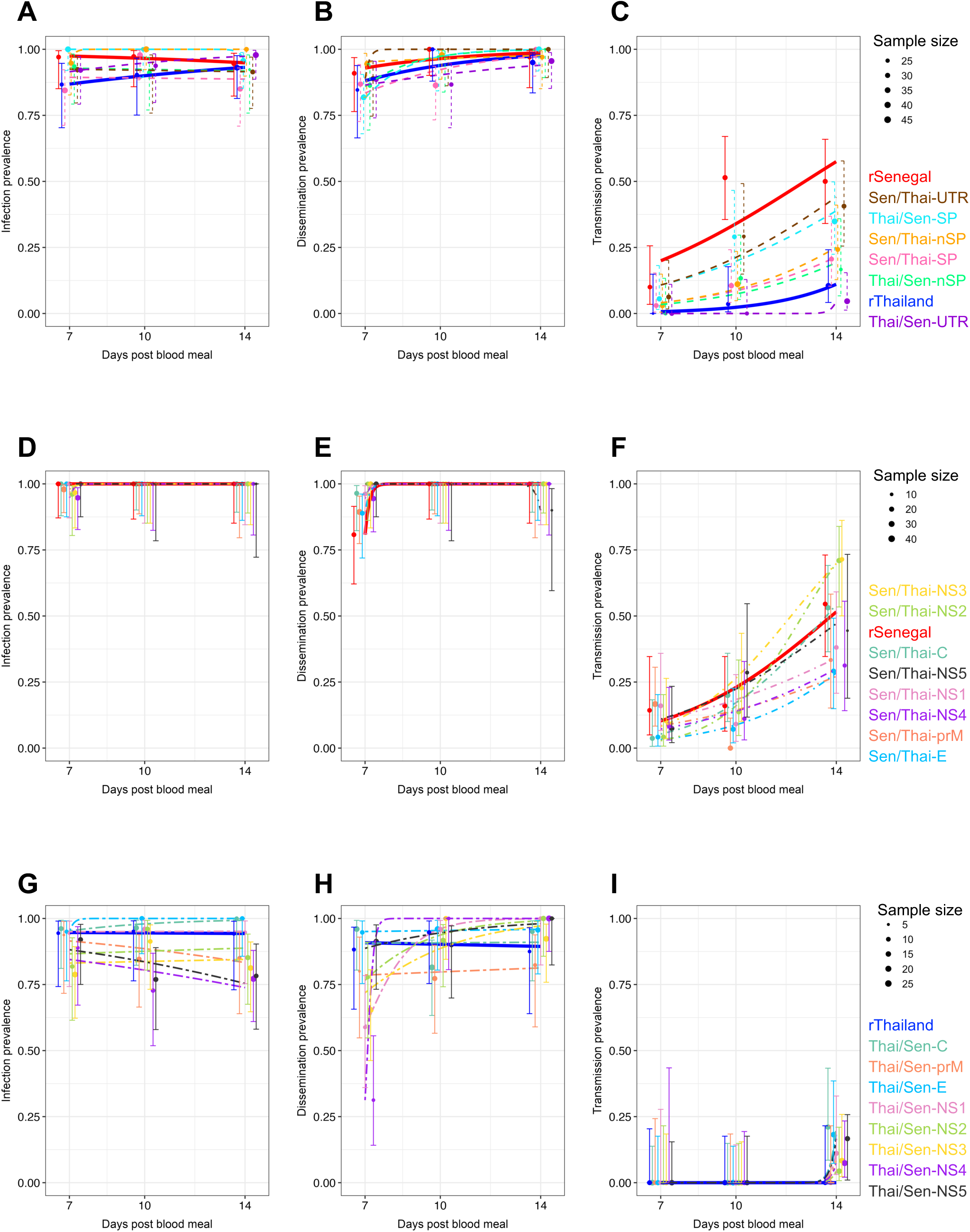
Structural and non-structural genes collectively drive differences in mosquito transmission between rSenegal and rThailand strains. *Aedes aegypti* mosquitoes from Colombia were orally exposed to the first set (**A**-**C**), second set (**D**-**F**), or third set (**G**-**I**) of chimeric viruses via an infectious blood meal. Mosquitoes were collected on days 7, 10, and 14 post infectious blood meal to assess the prevalence of midgut infection (**A**, **D**, **G**), systemic viral dissemination (**B**, **E**, **H**), and transmission potential (**C**, **F**, **I**) for each ZIKV strain. Infection prevalence is the proportion of blood-fed mosquitoes with a virus-positive body (measured by RT-PCR). Dissemination prevalence is the proportion of infected mosquitoes with a virus-positive head (measured by RT-PCR). Transmission prevalence is the proportion of mosquitoes with a disseminated infection and infectious saliva (measured by focus-forming assay). (**A**-**I**) Data points show the empirically measured proportions, with point size proportional to sample size (number of mosquitoes). Logistic regression results are represented by fitted lines, with error bars indicating 95% confidence intervals for the logistic fits. Lines and points are color-coded to represent the different chimeric viruses, and the parental strains are depicted with thicker lines. Raw data and logistic regression results are provided in Tables S3 and S4, respectively.

### Individual genes have a limited impact on viral growth in mosquito cells and mosquito transmissibility

To narrow down the genome regions responsible for differences in mosquito transmissibility between the rSenegal and rThailand strains, we constructed a second set of chimeric viruses by substituting each viral gene in the rSenegal strain with the corresponding gene from the rThailand strain (Figure S4A), and a third set by substituting each viral gene in the rThailand strain with the corresponding rSenegal gene (Figure S4B). Introducing specific genes such as *prM*, *E*, or *NS1* from the rThailand strain into the rSenegal strain resulted in smaller plaques on Vero E6 cells, while replacement of *NS1* and *NS4* led to larger plaques (Figure S4C). Conversely, all chimeric viruses of the third set formed smaller plaques than the parental rThailand strain, indicating an asymmetric influence on plaque size (Figure S4D). We then compared the growth kinetics of the new sets of chimeric viruses in mosquito cells with that of the parental strains. In the second set, all replacements significantly reduced infectious viral titers except *C* and *NS1* (Figure S4E). In the third set, replacement of *C*, *E*, or *NS5* transiently increased virus titers at 48 and 72 h.p.i. but resulted in lower titers than rThailand at 96 h.p.i., while replacing *NS2* led to a significantly slower growth (Figure S4F). Taken together, these results show that substitutions of *E* or *NS5* most significantly impact virus growth kinetics in mosquito cells, but no single-gene substitution significantly alters the efficiency of infectious particle production in both directions.

To assess how individual viral genes affect mosquito transmissibility, we exposed mosquitoes from Colombia to the second and third sets of chimeric viruses via artificial infectious blood meals containing 2×10^6^ PFU/ml (Figure S1B). Actual blood meal titers varied from 0.7 to 2.5×10^6^ PFU/ml and from 1.2 to 2.1×10^6^ PFU/ml for the second and third sets, respectively, and this variation was factored into our statistical analysis (Table S4). At least 10 and 16 mosquitoes per time point were tested for each virus for the second and third sets, respectively (Table S3). Virtually all mosquitoes exposed to the second set of chimeric viruses were infected and had a disseminated infection, with no significant variation among viruses (Figure 3D-E). Transmission prevalence increased over time, reaching 29%-71% on day 14 (Figure 3F), with a marginally significant effect of the virus (Table S4). Following exposure to the third set of chimeric viruses, both infection prevalence and dissemination prevalence were significantly influenced by the virus (Table S4), however the magnitude of these differences was modest, ranging on average from 78.7% to 98.5% and from 79.6% to 95.5%, respectively (Figure 3G-H). Transmission prevalence significantly increased over time but was not influenced by the virus (Figure 3I; Table S4). Together, these results show that no single viral gene is solely responsible for the observed variation in transmission prevalence in mosquitoes from Colombia.

### Structural genes influence virus transmission irrespective of mosquito genetic background

To investigate if the observed differences in transmission prevalence between chimeric viruses (Figure 3C) were specific to the mosquitoes from Colombia, we tested the first set of chimeric viruses in two other mosquito colonies with different genetic backgrounds and contrasting levels of ZIKV susceptibility. ZIKV susceptibility is known to be significantly higher in globally invasive populations of *Ae. aegypti* found outside Africa than in native African populations (*47*). We chose a ZIKV-resistant mosquito colony from Uganda representative of the native African populations (*47*), and a colony from Cape Verde with admixed genetic ancestry and an intermediate level of ZIKV susceptibility (*17*). We challenged mosquitoes from Uganda and Cape Verde with artificial infectious blood meals containing 1×10^7^ PFU/ml and 2×10^6^ PFU/ml, respectively, to maximize infection prevalence in these relatively resistant colonies. Actual blood meal titers varied from 3.0 to 8.0×10^6^ PFU/ml and from 0.7 to 4.0×10^6^ PFU/ml for the mosquitoes from Uganda and Cape Verde, respectively, and this variation was factored into our statistical analysis (Table S5). In both mosquito colonies, the virus significantly influenced transmission efficiency (Table S5). The rSenegal strain resulted in significantly higher transmission efficiency than the rThailand strain and swapping either SPs or nSPs—but not UTRs—between the parental strains resulted in intermediate transmission efficiencies that were consistent with the patterns previously observed with the mosquitoes from Colombia (Figure S5). Overall, the influence of SPs on transmission efficiency was more pronounced than the influence of nSPs. These results indicate that the viral genetic determinants of ZIKV transmission efficiency are largely independent of the mosquito genetic background.

### Relatively simple models explain viral growth *in vitro* and the dose response of midgut infection

To understand the mechanisms behind the varying mosquito transmissibility of chimeric viruses in mosquitoes, we developed a stochastic model of *in vivo* viral dynamics that could reproduce the qualitative outcomes of oral exposure to the chimeric viruses, capturing patterns of infection, dissemination, and transmission. Following an infectious blood meal, the virus first infects and replicates within the midgut epithelial cells, then ‘escapes’ from the midgut and disseminates via the hemocoel to other tissues. It ultimately reaches the salivary glands, where it is released in the saliva and transmitted to the next host (*48–50*). We aimed to use the stochastic model to pinpoint the key processes within mosquitoes that are most likely influenced by differences among the chimeric viruses. The output of our logistic model of *in vitro* viral growth kinetics (Figure 4A) was consistent with our time-course analyses of infectious particle production using the first set of chimeric viruses when the maximum titer supported by the cell culture (carrying capacity, *k*) was fixed at 10^8.5^ PFU/ml, the starting virus concentration (*s*) was either 10^1^ or 10^1.3^ PFU/ml, and the growth rate (*r*) was either 0.035 or 0.05 h^-1^, respectively (Figure S6A). We applied this viral growth model to capture *in vivo* mosquito infection dynamics, incorporating additional parameters: the probability of midgut infection (*β*), representing the likelihood that the virus successfully infects the midgut; the escape rate (*λ*), denoting how quickly the virus transfers between different tissues; and the blood meal clearance rate (*μ*), indicating how rapidly the virus is lost within the ingested blood (Figure 4B). Our sensitivity analyses showed that within the ranges of the parameter values used, the probability of midgut infection (*β*) had the strongest effect on the proportion of simulations with midgut infection, followed by the blood meal clearance rate (*μ*) (Figure S7). In scenarios where the starting virus concentration was 10^6^ PFU/ml, all simulations resulted in successful infections when *β* ranged from 10^-4.7^ to 10^-4^ and *μ* varied between 0.014 and 0.08 h^-1^. We subsequently used these parameter values in sensitivity analyses to evaluate viral dissemination and potential transmission (Figure S8). We found that by using values of *β* between 10^-5^ and 10^-3^ (Figure S6B), we could simulate the dose-response curves of midgut infection by the first set of chimeric viruses (Figure S3). These results suggest that differences in the dose-response curves of midgut infection across the chimeric viruses likely reflect variations in *β*, encompassing all processes from viral attachment, internalization, and replication in midgut cells.

**Figure 4.**
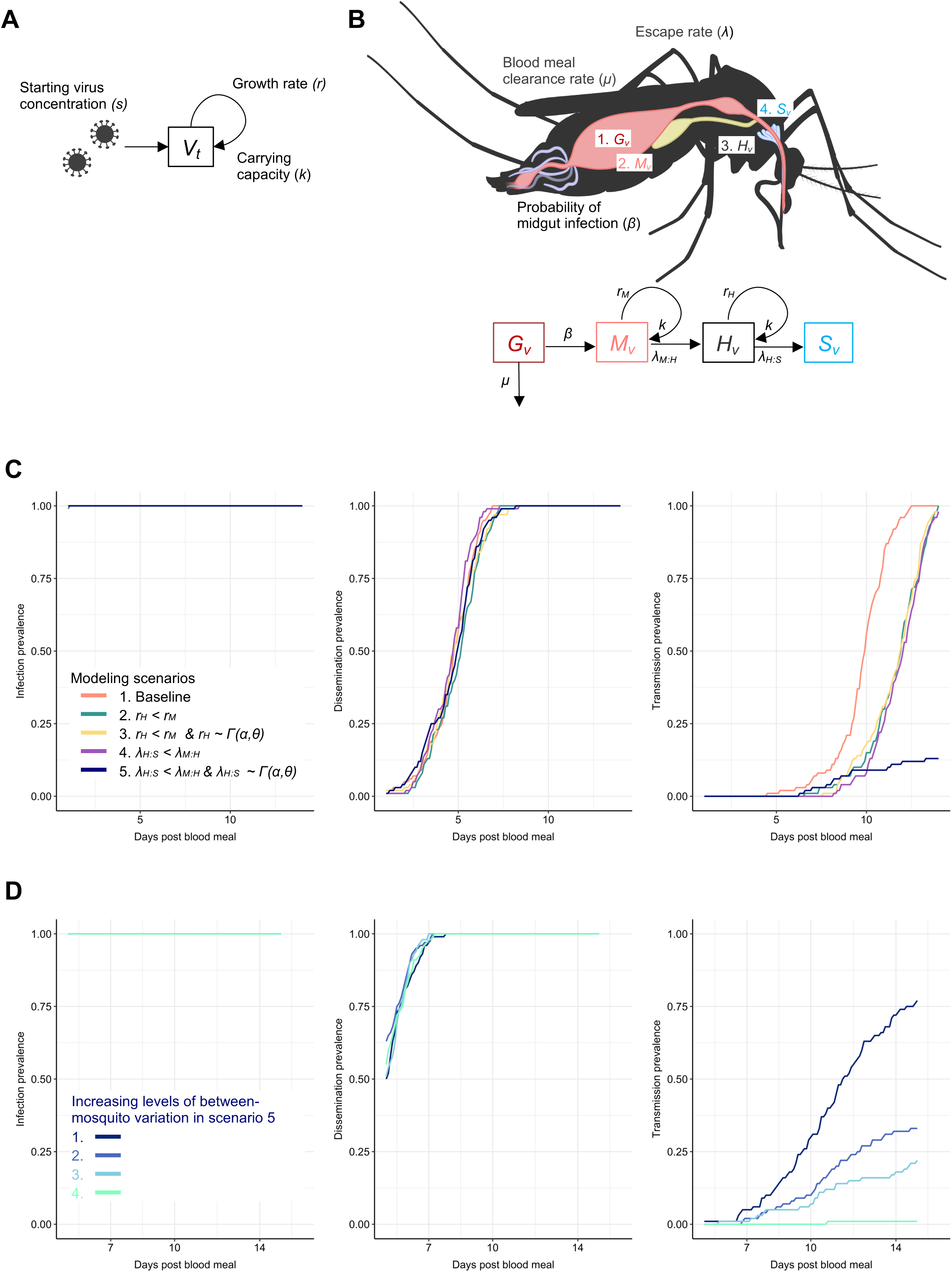
Modeling ZIKV infection dynamics identifies salivary gland traversal as the primary driver of differences in transmissibility. (**A**) Conceptual model for *in vitro* viral dynamics. The dynamics are described using a logistic growth curve as per Eq. 1 (see Methods). (**B**) Conceptual model for *in vivo* viral dynamics. The mosquito image is from BioRender.com. After ingestion of a blood meal containing infectious virus (*G_v_*), the virions are degraded in the blood meal according to a clearance rate (*μ*). The probability that at least one virion infects the midgut epithelium (*β*) determines whether infection is established. If infection occurs in the midgut (*M_v_*), the virus replicates at a growth rate (*r*) constrained by the carrying capacity (*k*). The virus may then disseminate to the hemocoel according to an ‘escape’ rate (*λ*). Virus in the hemocoel (*H_v_*) undergoes similar replication dynamics as in the midgut and can ‘escape’ to infect the salivary glands (*S_v_*), which eventually enables virus release into saliva. The simplest model assumes fixed parameter values (*r*, *k*, *λ*) across tissues and no between-mosquito variation in probabilities or rates. These assumptions are relaxed stepwise to evaluate processes underlying experimental observations. (**C**) Model outputs for viral dissemination. The proportion of simulations reaching the hemocoel is shown for five model scenarios (Table 2). Across all scenarios, except scenario five, transmission occurs in 100% of simulations. Scenario five introduces random variation in the transfer of virions between the hemocoel and salivary glands, enabling a reduced proportion of simulations with transmission, consistent with experimental findings. (**D**) Proposed hypothesis for difference between chimeric viruses of the first set. Results from scenario five suggest that differences in chimeric viruses may arise from variability in the rate at which virions infect the salivary glands and/or are released into saliva. This variability can be represented by a Gamma distribution, with variance adjusted between simulations to reflect distinct virus-mosquito interactions. Example Gamma distributions were modeled with variances of 10⁻⁷.⁵, 10⁻⁷.³, 10⁻⁷, and 10⁻⁶.⁵ to show the effects of this variation.

### Observed differences in transmission prevalence can be explained by between-mosquito variation in salivary gland infection

When we exposed mosquitoes from Colombia to the first set of chimeric viruses, despite >90% dissemination prevalence across all viruses by day 14 (Figure 3B), we observed lower transmission prevalence overall (<50%) and significant variation between viruses (Figure 3C). Our sensitivity analysis showed that the growth rate (*r*) was the most influential parameter, at least within the ranges of parameter values used, on viral dissemination in the hemocoel on day seven (Figure S8A) and transmission potential (salivary gland infection) on day 10 post infectious blood meal (Figure S8B). There was a relatively narrow range of the growth rate (*r*) values (between 0.05 and 0.1 h^-1^) where the proportion of simulations went from zero to one for dissemination on day 7, and for transmission potential on day 10. Our analysis of potential causes for the observed differences in transmission dynamics between chimeric viruses focused on variations in *r* and *λ* between tissues. We explored alternative model parameterizations for *r* and *λ* by using tissue-specific parameter values and by introducing Gamma distributions to account for between-mosquito variation. Of the five hypothesized scenarios (Table 2), only one—featuring a lower hemocoel-to-salivary gland escape rate (*λ_H:S_*) compared with the midgut-to-hemocoel escape rate (*λ_M:H_*) and between-mosquito variation in this parameter—resulted in 100% prevalence of infection and dissemination but less than 60% transmission prevalence (Figure 4C and scenario five, Table 2). The other scenarios primarily shifted the transmission prevalence curve to the right. Manipulating the variance of the Gamma distribution of *λ_H:S_* reproduced the differences in transmission prevalence observed between the chimeric viruses (Figure 4D), supporting the conclusion that these differences likely reflect variation in the hemocoel-to-salivary gland escape rate.

## Discussion

Previous experimental infections of mosquitoes with ZIKV strains from the African and Asian lineages have demonstrated significant variability in mosquito-borne transmission efficiency (*27*). However, the genetic determinants underlying these differences remain poorly understood. To address this gap, we developed chimeric viruses derived from recent African and Asian ZIKV parental strains, incorporating the numerous amino-acid substitutions and silent mutations distinguishing the parental genomes. Our findings revealed that African SPs critically enhance viral internalization, while African nSPs significantly improve viral genome replication in mosquito cells *in vitro*, relative to their Asian counterparts. Furthermore, *in vivo* experiments with these chimeric viruses demonstrated that differences in mosquito-borne transmissibility between African and Asian ZIKV are primarily determined by the SPs and, to a lesser extent, the nSPs. Substitutions in SPs such as E or prM—known to form essential heterodimers for maturing infectious virus particles (*51, 52*)—markedly enhanced replication *in vitro*, and increased transmission prevalence *in vivo* when introduced from the African to the Asian strain. The consistency of our results across *Ae. aegypti* populations with varying levels of ZIKV susceptibility highlights the robustness of SPs effects on transmission efficiency and their independence from population-specific mosquito factors. We found that UTRs had no significant effect on viral growth *in vitro* or transmission efficiency *in vivo*, a finding that contrasts with previous studies. Despite the proposed role of non-coding subgenomic flavivirus RNAs derived from the 3’-UTR in mosquito-borne transmission of ZIKV (*53*) and other orthoflaviviruses (*54–56*), lineage-specific differences in the ZIKV UTRs did not contribute to variation in mosquito transmissibility in our study.

In general, the chimeric viruses showed intermediate phenotypes, remaining within the range defined by the parental strains. Both SPs and nSPs were necessary to replicate the parental traits, and we found that cumulative effects of multiple genes, rather than any single gene, shape ZIKV transmissibility. These findings suggest that ZIKV transmissibility is governed by complex interactions among multiple viral genes, which act across various stages of virus propagation within mosquito cells. Future studies incorporating targeted combinations of viral gene substitutions, guided by these results, could further elucidate the genetic basis of transmissibility. The observed similarity between engineered and natural ZIKV strains establishes the validity of our reverse genetics system for studying mosquito transmissibility. Such precision in recapitulating phenotypes enables controlled dissection of genetic factors underlying epidemiologically relevant phenotypic differences.

Experimental infections of mosquitoes with viruses often provide only static snapshots of the infection process, typically focusing on individual stages of propagation, or fixed time points post exposure. This limited scope makes it challenging to understand more fully the dynamic processes governing viral propagation in mosquitoes. In this study, we addressed this complexity by integrating experimental data from chimeric viruses into a mathematical model. Consistent with experimental findings, the stochastic model pointed to the hemocoel-to-salivary gland transition as the primary determinant of transmission efficiency in infected mosquitoes. Introducing between-mosquito variability in this transition reproduced differences in transmission prevalence across chimeric viruses, supporting individual-level heterogeneity as a key factor. Together, these results indicate that once a mosquito is infected, ZIKV transmission predominantly depends on the ability of the virus to traverse the mosquito salivary glands, likely involving an infection barrier and/or an escape barrier to viral release into saliva. Unlike midgut infection, which requires the virus to navigate from the apical to the basal side of epithelial cells, secondary infection of salivary glands operates through a distinct mechanism that results in virus release in salivary secretions. Research on the vesicular stomatitis virus glycoprotein G in *Drosophila melanogaster* suggests that a specific signal motif in the viral envelope protein is critical for viral movement within midgut cells but less so in salivary gland cells, emphasizing the need to investigate the role of ZIKV SPs in polarized trafficking and virus release into saliva (*57*). Additionally, the dissemination of ZIKV from the hemocoel to the salivary glands warrants further exploration. Hemocytes, known for their antiviral immune roles in mosquitoes, have been observed to assist ZIKV spread to the ovaries and salivary glands after infection (*50*). Future research should focus on tissue-specific mechanisms of virus proliferation, including interactions within the hemolymph and salivary glands.

It is worth noting that most of our experiments used a standardized infectious dose in the artificial blood meals that infected 80-100% of the mosquitoes across the chimeric viruses. However, our initial dose-response experiment showed that chimeric viruses also differed in the ability to infect mosquitoes at lower doses (Figure S3; Table S3). Both SPs and nSPs from the African strain contributed to its higher infectivity in mosquitoes compared to the Asian strain. A previous study using *in situ* immunofluorescence showed that an African ZIKV strain established midgut infections and replicated more efficiently in midgut epithelial cells than an Asian strain (*58*). Therefore, it is likely that both superior midgut infectivity and more efficient traversal of the salivary glands contribute to higher transmissibility of African ZIKV strains in a real-world situation.

The absence of recorded outbreaks caused by ZIKV strains of the African lineage, despite their higher transmissibility, presents a paradox. The lower ZIKV susceptibility of African *Ae. aegypti* populations may have played a role in preventing ZIKV outbreaks on the continent (*47*). Alternatively, the lower mosquito transmissibility of Asian ZIKV strains might be compensated by a potential fitness advantage within human hosts. The epidemiological fitness of ZIKV depends not only on the efficiency of mosquito-borne transmission but also on the level of human infectiousness to mosquitoes. In mouse models, African ZIKV strains typically exhibit higher viremia peaks compared to Asian strains (*27*), but the relative magnitude and duration of infectiousness to mosquitoes remain unclear. Theoretical models indicate that while high viremia may lead to a rapid decline in titer, a lower yet more sustained viremia could be more advantageous for viral transmission (*59*). Future research should investigate whether a more gradual ZIKV viremia in humans correlates with a prolonged period of infectiousness to mosquitoes (*60*).

Epidemic strains of mosquito-borne viruses frequently harbor mutations that facilitate human transmission and consequently, intensify outbreaks (*61*). Previous studies identified mutations that enhance the replication of Asian lineage ZIKV strains in *Ae. aegypti* mosquitoes, which arose just prior to ZIKV emergence in the Americas (*36, 37*). These transmission-adaptive variants are thought to be ancestral because they are also found in African lineage ZIKV strains (*38*). However, individually, these mutations do not recapitulate the level of transmissibility observed in African lineage ZIKV strains (*38*). Our findings indicate that the greater transmissibility of African lineage ZIKV strains compared to their Asian counterparts involves multiple genetic factors rather than single mutations. Thus, the polygenic nature of ZIKV transmissibility may have prevented Asian lineage strains from achieving the same epidemic potential as African lineage strains. These results underscore the complexity of viral adaptation, which is often limited by epistatic interactions and lineage-specific adaptive landscapes (*62*). Elucidating the complex viral genetic basis of mosquito-borne transmission is critical for understanding viral evolution and improving our ability to predict and effectively manage epidemic situations.

## Materials and methods

### Ethics and regulatory information

All experiments were conducted in compliance with national guidelines and with European Commission Directive 2000/54/EC on the protection of workers from risks related to exposure to biological agents at work, and with Directives 2009/41/EC and 98/81/EC on the contained use of genetically modified micro-organisms. The generation of chimeric viruses and their use in experimental infections of mosquitoes were approved by the French Ministry of Higher Education, Research, and Technology (authorization number 8933) and the Institut Pasteur Dual Use Liaison Group (DURC 2021-03). Wild mosquito eggs were collected and exported with permission from local institutions and/or governments as required (Uganda: permit 2014-12-134; Cape Verde: authorization No 988).

### Cells

Huh-7 and Vero E6 cells were maintained at 37°C under 5% CO_2_ in high-glucose Dulbecco’s modified Eagle’s medium (DMEM) containing 10% fetal bovine serum (FBS; Gibco Thermo Fisher Scientific) and 1% penicillin/streptomycin (pen/strep; Gibco Thermo Fisher Scientific). C6/36 cells were maintained at 28°C in Leibovitz’s L-15 medium (L15; Gibco Thermo Fisher Scientific) containing 10% FBS, 2% tryptose phosphate broth (TPB; Gibco Thermo Fisher Scientific), 1× non-essential amino acids (NEAA; Gibco Thermo Fisher Scientific), and 1% pen/strep.

### Wild-type viruses

Wild-type ZIKV strain Kedougou2011, referred to as the iSenegal strain in this study, was isolated near Kédougou in 2011 from a pool of wild-caught mosquitoes (*63*). Wild-type ZIKV strain THA/2014/SV0127-14, referred to as the iThailand strain in this study, was isolated from a human serum sample (*64*). Virus stocks were prepared in C6/36 cells and the viral genome sequences were obtained by high-throughput sequencing as previously described (*27, 65*). Wild-type ZIKV strains iSenegal and iThailand were converted by reverse genetics into rSenegal and rThailand strains, respectively, as described below for chimeric viruses.

### Chimeric viruses

Parental and chimeric ZIKV strains were generated by circular polymerase extension reaction (CPER) following a published method (*66*) with modifications. For each virus, a total of six ZIKV complementary DNA (cDNA) fragments covering the full-length viral genome were amplified utilizing PrimeSTAR GXL DNA Polymerase (TaKaRa Bio) and cloned into the pMW118 vector (listed in the Reagent Table). A CMV linker for ZIKV was then inserted into the pCR-Blunt II-TOPO vector, encoding sequences of the HDVr, Late SV40 pA signal, and CMV promoter. All DNA inserts were verified through Sanger sequencing. Following the cloning process (with some optimizations shown in Figure S1A), infectious cDNA was generated by assembling DNA fragments F1-F6, which encompass the entire ZIKV genome, and fragment F7, encoding HDVr, SV40 pA signal, and CMV promoter. The fragments were amplified, and the chimeric viral genomes were introduced using cloning plasmids as templates along with specific primer sets detailed in Tables S1 and S2. All the DNA fragments were designed to have complementary ends with a 49- to 95-nucleotide overlap. Equimolar amounts (0.1 pmol each) of the resulting DNA fragments F1-F7 were mixed in 50-μl reaction volumes of PrimeStar GXL with 2 μl of DNA polymerase. CPER was carried out with an initial 2 minutes of denaturation at 98°C; 20 cycles of 10 seconds at 98°C, 15 seconds at 55°C, and 12 minutes at 68°C; and a final extension for 12 minutes at 68°C. The resulting CPER products, encoding the CMV promoter, full-length ZIKV genome sequence, followed by HDVr and SV40 poly(A) signal, were directly transfected into Huh-7 cells using Trans IT LT-1 (Mirus) following the manufacturer’s protocol. The culture supernatants from Huh-7 cells were collected and inoculated onto C6/36 cells. After three passages in C6/36 cells, the virus stocks were centrifuged to remove cell debris and stored at -80°C for future use. Virus stock titers in plaque-forming units/ml (PFU/ml) were determined by plaque assay and full-genome sequences were confirmed by high-throughput sequencing as described below.

### Viral genome sequencing

The full-length sequence of parental and chimeric ZIKV genomes was determined by Illumina sequencing as previously described (*67*). Briefly, total RNA was extracted from a virus stock using QIAamp Viral RNA Mini Kit (Qiagen) and treated with Turbo DNase (Invitrogen). After the RNA purification with RNAClean XP beads (Beckman Coulter), cDNA was synthesized using M-MLV Reverse Transcriptase (Invitrogen), random hexameric primers (Roche) and RNaseOUT Recombinant Ribonuclease Inhibitor (Thermo Fisher Scientific) according to the manufacturer’s protocol. Double-stranded DNA (dsDNA) was produced with Second-Strand Synthesis Buffer (New England Biolabs), *E. coli* DNA ligase (New England Biolabs), *E. coli* DNA polymerase I (New England Biolabs) and *E. coli* RNase H (New England Biolabs), followed by DNA purification with AMPure XP beads (Beckman Coulter). The dsDNA was quantified by Qubit dsDNA HS assay kit (Invitrogen) and used for library preparation with a Nextera XT Library preparation kit (Illumina) according to the manufacturer’s instructions. The final libraries were checked with Bioanalyzer high sensitivity DNA analysis (Agilent) and sequenced on an Ilumina NextSeq 500 instrument (150 cycles, paired ends) with NextSeq 500/550 v2.0 Kit (Illumina). Adapters and low-quality sequences of raw reads were removed using Trimmomatic v0.39 (*68*). The trimmed reads were assembled using megahit v1.2.9 (*69*) with default parameters. The contigs were queried against the NCBI non-redundant protein database using DIAMOND v2.0.4 (*70*), to look for potential contaminants in addition to the detected ZIKV genome. ZIKV scaffolds were constructed using the longest assembled contig and a viral sequence obtained in the previous study (*27*). The scaffolds were used to map the trimmed reads, using clc-assembly-cell v5.1.0. The consensus sequence generation and intrasample variant analysis were performed with ivar v1.0 (*71*) using a minimum of 5× read depth of coverage for the consensus, and 500× with a 2% minimum threshold for minor variants. Samtools v1.10 (*72*) was used to sort the aligned BAM files and generate alignment statistics. The mapping data was visually checked to confirm the accuracy of the obtained genomes using Geneious Prime 2023 (www.geneious.com). The raw reads were deposited in the Sequence Read Archive under bioproject PRJNA1199883, and the full-length consensus genome sequences of the viruses were deposited in GenBank under accession numbers PQ869243-PQ869266.

### Plaque assay

Vero E6 cells were seeded in 24-well plates one day prior to virus inoculation. Ten-fold serial dilutions of the samples were prepared in DMEM and inoculated onto the confluent Vero E6 cells after removing the cell-culture supernatant. After 1-hour incubation at 37°C, the inoculum was replaced by DMEM containing 1.0% carboxymethylcellulose (Avantor, VWR), 1% FBS, and 1% pen/strep and the cells were incubated for 7 days at 37°C. To count plaque numbers, the cells were fixed with 4% formaldehyde solution (Sigma) and stained with 0.2% crystal violet (Sigma).

### Viral growth kinetics *in vitro*

C6/36 cells were seeded in 6-well plates one day prior to virus inoculation. After removing the cell-culture supernatant, the confluent cells were inoculated with ZIKV at a multiplicity of infection (MOI) of 0.01 for 1 hour. The virus inoculum was replaced by fresh L15 medium containing 2% FBS, TPB, NEAA, and 1% pen/strep (2% FBS-L15 medium) and the cells were incubated at 28°C. At 24, 48, 72, and 96 hours post infection (h.p.i.), culture supernatants were collected, and infectious titers were determined by plaque assay as described above.

### Virus attachment assay

C6/36 cells were seeded in 24-well plates one day before virus inoculation. The confluent cells were pre-chilled on ice for 15 minutes before the start of the assay. After removing the cell-culture supernatant, ZIKV was added to the cells at an MOI of 1 to allow virus attachment. After 1-hour incubation on ice, the supernatant was collected for RNA extraction with QIAamp Viral RNA Mini Kit (Qiagen). The cells were rinsed twice with pre-chilled PBS before being lysed using TRIzol (Invitrogen). Total RNA was then extracted following the manufacturer’s protocol. Viral RNA was quantified by quantitative reverse transcription PCR (RT-qPCR) for ZIKV on total RNA using GoTaq Probe 1-step RT-qPCR kit (Promega), following the manufacturer’s protocol. The primers, probes, and gBlocks utilized in the RT-qPCR for ZIKV are listed in Table S2.

### Virus internalization assay

C6/36 cells were seeded in 24-well plates and pre-chilled on ice for 15 minutes before the start of the assay. As controls, cells were pre-treated with Dynasore (Sigma) at a final concentration of 100 uM in 2% FBS-L15 medium or with an equivalent dilution of DMSO for 1 hour before chilling. After removing the cell-culture supernatant, ZIKV was added to the cells at an MOI of 1 and incubated on ice for 1 hour to allow virus attachment. The cells were washed with PBS to remove unbound virus particles, and fresh 2% FBS-L15 medium containing the same concentrations of Dynasore or DMSO was added. The virus was then allowed to internalize into the cells for 3 hours at 28°C. After this period, the cells were washed with PBS and treated with protease E (5 mg/ml, Sigma) for 15 minutes on ice to remove any viruses remaining on the cell surface. The cells were washed 3 times with PBS and subsequently lysed with TRIzol (Invitrogen) for RNA extraction. The levels of viral RNA were quantified in total RNA by RT-qPCR using GoTaq 1-step RT-qPCR kit (Promega). *Actin* expression was assessed as an internal control using the same RT-qPCR method and the primer set listed in Table S2. The efficiency of virus internalization, measured at 3 h.p.i, was determined by calculating the ratio of intracellular viral RNA to *Actin* expression. This ratio was normalized to the ratio of viral RNA to *Actin* expression from cells sampled at 0 h.p.i. without protease E treatment.

### Quantification of ZIKV genomic and antigenomic RNA

C6/36 cells were seeded in 24-well plates and pre-chilled on ice for 15 minutes before the start of the assay. The confluent cells were inoculated with ZIKV at an MOI of 1 for one hour on ice to allow virus attachment. After one hour, the cells were washed with PBS and fresh 2% FBS-L15 medium was added. The cells were then incubated at 28°C, and samples were collected at 0, 3, 6, 9, 12, 15, 18, 21, and 24 h.p.i. Each time point involved washing the cells with PBS, treating them with 5 mg/ml protease E for 15 minutes on ice, and lysing in TRIzol for total RNA extraction. Viral genomic and antigenomic RNAs were quantified using strand-specific RT-qPCR as previously described (*73, 74*), with minor optimizations. The standard curve for antigenomic RNA was constructed by amplifying qPCR targets using primers listed in Table S2, followed by reverse transcription using the MEGAscript T7 Transcription kit (Invitrogen) and purification using the MEGAclear Transcription Clean-Up kit (Invitrogen) as per the manufacturer’s guidelines. RNA concentrations were measured using a Nanodrop spectrophotometer, and 10-fold serial dilutions were made for standard curves. For specific incorporation of the tag sequence into cDNA, total RNA extracted from ZIKV-infected cells was reverse transcribed using M-MLV Reverse Transcriptase (Invitrogen) with RT-specific primers (Table S3). Quantification of cDNA was then performed using the GoTaq probe qPCR Master Mix (Promega) and a set of strand-specific and tag-specific primers listed in Table S2. At each time point, the ratio of viral antigenomic or genomic RNA to *Actin* expression was calculated as described above. To assess replication efficiency, relative viral antigenomic RNA was normalized against the ratio at 12 h.p.i., and relative genomic RNA against the ratio at 0 h.p.i.

### Infectious titer/viral genome ratio

C6/36 cells were seeded in 24-well plates one day prior to virus inoculation. The confluent cells were inoculated with ZIKV at an MOI of 1 for one hour at 28°C. After one hour, the inoculum was replaced by fresh 2% FBS-L15 medium. Cell-culture supernatant was sampled at 24, 48, 72, and 96 h.p.i. to determine the infectious titer by plaque assay and the concentration of viral genomic RNA by RT-qPCR, as described above.

### Virus decay

ZIKV was prepared at a titer of 10^6^ PFU/ml and incubated at 28°C for 96 hours. Samples were collected after 0, 6, 12, 24, 48, 72, and 96 hours, to determine infectious titer by plaque assay, as described above.

### Cell viability assay

C6/36 cells were seeded in 24-well plates one day prior to virus inoculation. The confluent cells were inoculated with ZIKV at a MOI of 0.01 for 1 hour at 28°C. After the 1-hour incubation, the inoculum was replaced by fresh 2% FBS-L15 medium. At the designated time points, the cell-culture supernatant was removed and CellTiter-Glo reagent (Promega) was added to the cells. Following a 10-minute incubation at room temperature (20-25°C), luminescence was measured using a GloMax 96 microplate luminometer (Promega) to quantify ATP, which is indicative of cell viability. For each viral infection, the cellular ATP levels at the indicated time points were normalized to the ATP levels of cells sampled at 0 h.p.i..

### Mosquitoes

*Ae. aegypti* colonies were originally established from wild specimens caught in Colombia in 2017, Uganda in 2015, and Cape Verde in 2020, as previously described (*17, 47*). Mosquitoes were reared under controlled insectary conditions (28° ± 1°C, 12-hour light/dark cycle and 70% relative humidity). Prior to performing the experiments, their eggs were hatched synchronously in a vacuum chamber for one hour. Larvae were reared in dechlorinated tap water supplemented with a standard diet of TetraMin fish food (Tetra). Adults were kept in 30×30×30-cm BugDorm-1 insect cages (BugDorm) with permanent access to 10% sucrose solution. Mosquito experimental infections were performed with the 16^th^-19^th^, 23^rd^, and 9^th^ laboratory generations of the colonies from Colombia, Uganda, and Cape Verde, respectively.

### Mosquito oral exposure to ZIKV

Mosquitoes were orally exposed to ZIKV by membrane feeding in a biosafety level-3 containment facility. Briefly, 5- to 7-day-old female mosquitoes were starved for one day prior to the oral challenge. The infectious blood meal comprised a 2:1 mixture of washed rabbit erythrocytes (BCL) and ZIKV suspension, supplemented with 10 mM adenosine triphosphate (Sigma) and 0.1% sodium bicarbonate (Sigma). Mosquitoes were allowed to feed on the infectious blood meal for 15 minutes via a membrane-feeding apparatus (Hemotek Ltd.) with porcine intestine serving as the membrane. After feeding, fully engorged females were sorted on ice, transferred to 1-pint cardboard containers, and maintained under controlled conditions (28° ± 1°C, 12-hour light/dark cycle with 70% relative humidity) within a climatic chamber, with permanent access to 10% sucrose solution. The infectious titer of the blood meal was verified by plaque assay as described above.

### Salivation assay

Mosquitoes were paralyzed with triethylamine (Sigma) for 5 minutes at 7, 10, and 14 days post infectious blood meal to collect saliva *in vitro* as previously described (*27*). Briefly, after removal of all legs, each mosquito’s proboscis was inserted into a 20-μl pipet tip containing 10 μl of FBS. The mosquitoes were allowed to salivate into this medium for 30 minutes. The saliva-containing FBS was then collected, combined with 40 μl of 2% FBS-DMEM supplemented with 4% Antibiotic-Antimycotic 100× (Life Technologies), and stored at −80°C for subsequent titration by focus-forming assay, as described below. After salivation, the heads and bodies of each mosquito were dissected and individually transferred to 300 μl of squash buffer, composed of 10 mM Tris pH 8.0, 50 mM NaCl, and 1.27 mM EDTA pH 8.0 (all from Invitrogen), supplemented with 0.35 mg/ml proteinase K (Eurobio Scientific). Head and body samples were stored at −80°C for subsequent testing by RT-PCR, as described below.

### Focus-forming assay

Vero E6 cells were seeded in 96-well plates one day prior to virus inoculation. Serial 10-fold dilutions of the samples were prepared in DMEM (except for saliva samples that were used undiluted) and inoculated onto the confluent cells after removal of the cell-culture supernatant. Following an incubation period at 37°C for 1 hour, the inoculum was replaced with DMEM containing 1.0% carboxymethylcellulose, 1% FBS, 1% pen/strep, and 4% Antibiotic-Antimycotic 100×. The cells were further incubated for 5 days at 37°C before being fixed with a 4% formaldehyde solution. For immunostaining, the fixed cells were permeabilized with 0.1% Triton X-100 in PBS (Sigma) for 10 minutes, blocked with 1% bovine serum albumin (Sigma) in PBS, and then incubated with a 1:1,000 dilution of mouse anti-flavivirus group antigen monoclonal antibody clone D1-4G2-4-15 (Merck) in PBS for 1 hour at room temperature (20-25°C). Following three washes with PBS, the cells were incubated with Alexa Fluor 488-conjugated goat anti-mouse IgG (Life Technologies) at a 1:1,000 dilution in PBS for 1 hour at room temperature. Fluorescent foci were visualized using an EVOS FL fluorescence microscope (Thermo Fisher Scientific) equipped with appropriate barrier and excitation filters.

### Detection of viral RNA by qualitative RT-PCR

To assess the presence of viral RNA in mosquito heads and bodies, the samples were homogenized for 30 seconds at 6,000 rotations per minute in a Precellys 24 grinder (Bertin Technologies). A 100-μl aliquot of the homogenate was transferred to a PCR plate and subjected to crude RNA extraction by incubation for 5 minutes at 56°C followed by 10 minutes at 98°C. Total RNA was then used to synthesize cDNA using M-MLV Reverse Transcriptase, RNaseOUT Recombinant Ribonuclease Inhibitor, and random hexameric primers, following the manufacturer’s instructions. The resulting cDNA was amplified by PCR utilizing DreamTaq DNA polymerase (Thermo Fisher Scientific) with two sets of primers: pair 1 comprised ZIKV-PCR-F (5’-GTATGGAATGGAGATAAGGCCCA-3’) and ZIKV-PCR-R (5’-ACCAGCACTGCCATTGATGTGC-3’), and pair 2 consisting of ZIKV-PCR-F and ZIKV-PCR-R1 (5’-TCGTATTGCCAACCAGGCCAAAGC-3’). PCR cycling conditions were set as follows: an initial denaturation for 2 minutes at 95°C, followed by 35 cycles of 30 seconds at 95°C, 30 seconds at 55°C, and 90 seconds at 72°C, concluding with a final extension of 7 minutes at 72°C. The PCR products were subsequently visualized via electrophoresis on a 1.5% agarose gel.

### Mosquito dose-response curves

To estimate the 50% oral infectious dose (OID_50_) of mosquitoes, dose-response curves were generated by preparing infectious blood meals containing 10^4^, 10^5^, and 10^6^ PFU/ml of ZIKV. Mosquitoes were orally exposed to ZIKV as described above. At 3- and 7-days post blood feeding, whole mosquito bodies were individually homogenized in 300 µl of 2% FBS-L15 supplemented with 4% Antibiotic-Antimycotic 100×. From each homogenate, 150 µl were used for total RNA extraction using the NucleoSpin96 kit (Macherey-Nagel), following the manufacturer’s protocol. Viral RNA detection was performed by qualitative RT-PCR as described above. The remaining homogenates were stored at −80°C for subsequent titration by focus-forming assay, as described above. The OID_50_ estimate was calculated from the dose-response curves using the drc package in R v.4.2.2 (www.r-project.org).

### Statistical analyses

The prevalence of ZIKV infection, dissemination, and transmission in mosquitoes was analyzed by logistic regression as a function of experiment, ZIKV strain, and time. The initial statistical model included all their interactions, which were removed from the final model if their effect was non-significant (*P* >0.05). Time was considered a continuous variable. To account for small, uncontrolled differences in virus concentration, the log_10_-transformed blood meal titer was also included in the initial model as a covariate and removed from the final model if non-significant (*P* <0.05). For *in vitro* assays, statistical significance was determined by two-tailed Student’s *t-*test or one-way analysis of variance (ANOVA) followed by Dunnett’s test. Statistical analyses were performed in JMP v.14.0.0 (www.jmp.com) and R v.4.2.2 (www.r-project.org).

### Model of *in vitro* viral dynamics

A logistic growth curve was employed to establish a model for predicting infectious viral titer (*V_t_*) in cell culture at time *t* post infection defined as:

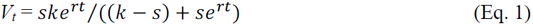

where *s* is the starting virus concentration, *r* is the growth rate, and *k is* the carrying capacity. Empirical data of viral growth kinetics in mosquito cells infected with the first set of chimeric viruses was used to approximate values for *s*, *r*, and *k*, enabling replication of dynamics observed in the laboratory experiments (Figure 2C). To simplify the modeling process and avoid complex statistical fitting, *s* and *k* were approximated based on the viral titer at the start and end of the experiments, respectively. The growth rate *r* was adjusted to ensure a reasonable match between model predictions and empirical growth curves.

### Model of *in vivo* viral dynamics

Following a previous study (*75*), the *in vitro* model (Eq. 1) was extended to simulate virus propagation within the mosquito midgut, hemocoel, and salivary glands. A probability of midgut infection (*β*) was incorporated to reflect the fact that only a few virions typically initiate infection out of thousands in the infectious blood meal (*76, 77*). This approach introduces a stochastic element into the model to reflect demographic variability observed in oral experimental infections of mosquitoes. The model entails six probabilistic processes (Table 1) affecting three key stages: i) initial probability (*β*) of midgut infection from virus in the blood meal, before viral clearance from the blood meal (*µ*); ii) viral replication (*r*), constrained by a carrying capacity (*k*) in the midgut and hemocoel; and iii) viral ‘escape’ (*λ*) from the midgut to the hemocoel (M:H) and from the hemocoel to the salivary glands (H:S). *G_v_*, *M_v_*, *H_v_*, and *S_v_* represent the virus levels in the blood meal, midgut, hemocoel, and salivary glands, respectively. The stochastic dynamics were simulated using the tau-leap version of the Gillespie algorithm (*78*). In the simplest model scenario (Table 2), the parameters for *r*, *k*, and *λ* were assumed to be the same across tissues.

**Table 1.** Transitions and reaction rates in the stochastic model of *in vivo* mosquito infection dynamics.

| Event | Change | Rate |
| --- | --- | --- |
| Blood meal clearance | $G_v - 1$ | $\mu G_v$ |
| Virus infection of the midgut | $G_v - 1, M_v + 1$ | $\beta G_v$ |
| Virus replication in the midgut | $M_v + 1$ | $r_M M_v (K - M_v) / K$ |
| Virus escape into the hemocoel (M:H) | $M_v - 1, H_v + 1$ | $\gamma_{MH} M_v$ |
| Virus replication in the hemocoel | $H_v + 1$ | $r_H H_v (K - H_v) / K$ |
| Virus invasion of the salivary glands (H:S) | $H_v - 1, S_v + 1$ | $\gamma_{HS} H_v$ |

**Table 2.** Model scenarios reflecting different hypotheses concerning how infection and dissemination processes may differ between mosquito tissues.

| Scenario | Model parameterization |
| --- | --- |
| 1. Baseline | All parameter values ($r$, $k$, $\lambda$) are the same between midgut and hemocoel tissues |
| 2. $r_H < r_M$ | Viral growth rate in the hemocoel ($r_H$) is less than the growth rate in the midgut ($r_M$) |
| 3. $r_H < r_M$ & $r_H \sim \Gamma(\alpha, \theta)$ | Viral growth rate in the hemocoel ($r_H$) is less than the growth rate in the midgut ($r_M$) and there is between-mosquito variation in $r_H$, such that for each simulation a value for $r_H$ is drawn randomly from a Gamma distribution with shape $\alpha$ and scale $\theta$. All other values are fixed across tissues. |
| 4. $\lambda_{H:S} < \lambda_{M:H}$ | The escape rate from the hemocoel to the salivary glands ($\lambda_{H:S}$) is less than the escape rate from the midgut to the hemocoel ($\lambda_{M:H}$). All other values are fixed across tissues. |
| 5. $\lambda_{H:S} < \lambda_{M:H}$ & $\lambda_{H:S} \sim \Gamma(\alpha, \theta)$ | The escape rate from the hemocoel to the salivary glands ($\lambda_{H:S}$) is less than the escape rate from the midgut to the hemocoel ($\lambda_{M:H}$). There is between-mosquito variation in $\lambda_{H:S}$, such that for each simulation a value for $\lambda_{H:S}$ is drawn randomly from a Gamma distribution with shape $\alpha$ and scale $\theta$. All other values are fixed across tissues. |

### Sensitivity analysis of the *in vivo* model

A sensitivity analysis was conducted to explore the effects of varying parameter values on the probability of midgut infection as a function of starting virus concentration, and of the probability of systemic dissemination and transmission as a function of time. Empirical dose-response curves were used to inform on ranges of parameter values that could potentially produce results qualitatively to match those observed in the experiments. The stochastic model was run 30 times for each subset of parameters to simulate viral dynamics in 30 individual mosquitoes. Midgut infection simulations ran for 124 hourly time steps, whereas dissemination and transmission simulations ran for 360 hourly time steps. For midgut infection, Latin hypercube sampling generated 1000 parameter sets within the following ranges: *μ* from 1/72 to 1/12 h^−1^; *β* from 0 to 0.0001; *r* from 0.001 to 0.1 h^−1^; and *k* from 10^3^ to 10^20^ PFU/ml. The proportion of mosquitoes developing a midgut infection was determined as the proportion of simulations where virus levels in the midgut exceeded zero (*M_v_* >0) at the end of the 124 hourly time steps. *G_v_* was set at 10^6^ PFU/ml, adjusted by multiplying by 0.003, reflecting the average mosquito blood meal volume (*79*). Using the same sampling approach, 1000 parameter sets were generated for viral dissemination (hemocoel) and transmission potential (salivary glands), adopting the maximal and minimal values of *μ* and *β* that resulted in successful midgut infections from earlier simulations. Parameter ranges for *r* and *k* remained unchanged. *λ* was set between 10^-8^ and 10^-5^ h^−1^. The outcome was determined as the proportion of simulations showing established infections in the hemocoel (*H_v_* ≥1), or salivary glands (*S_v_* ≥1), over intervals of 24 time steps (equivalent to per day), taking the last output for each daily time step. Evaluation threshold was set at 1000 PFU/ml to confirm infection establishment.

### Simulating the *in vivo* data with the model

Initially, the model was used to reproduce, qualitatively, the dose-response curves of ZIKV infection prevalence observed experimentally. By adjusting *β* of at least one virion infecting the midgut epithelium, the model was calibrated to match empirical observations with the first set of chimeric viruses. Each value of *β* was tested over 124 hourly time steps in 100 simulations across varying *G_v_* values from 10^2^ and 10^8^ PFU/ml to quantify the proportion of established midgut infections, as described above. Following the results from the sensitivity analyses, *μ* of 1/72 h^-1^, *r* of 0.04 h^-1^, *k* of 10^20^ PFU/ml, and *λ* of 0.00005 h^-1^ were kept constant. Next, parameter values of *β* derived from dose-response curves *in vivo* and of *r* estimated from growth kinetics in mosquito cells were combined to simulate virus dissemination to the hemocoel and salivary glands. Simulations aimed to explore five different scenarios to understand patterns of virus spread within mosquitoes and to provide biologically plausible hypotheses to explain the experimental data (Table 2). Of note, the process of salivary gland infection and virus escape into the saliva were collapsed into *S_v_* in the simulations, whereas the experiments detected virus presence in expectorated saliva. The baseline model (scenario 1, Table 2) was used as a simplistic framework where uniform infection and growth parameters were assumed to enable comparisons against four more elaborate scenarios. Differences in the infection processes are expected, such as variations in *r* across different cell types (*80*), cell-to-cell viral spread within the midgut, and virus propagation via freely moving hemocytes in the hemocoel (*49, 50*), and distinct anatomical barriers between the midgut, hemocoel, and salivary glands (*81, 82*). Additional model parameterizations were explored to represent the observed tissue-specific infection patterns more accurately. These modifications were aimed at modeling widespread infection across simulations consistently, though not necessarily achieving transmission in every single mosquito or each simulation (Figure 4). The four more complex scenarios were designed to reflect variations in processes between tissues that could be consistent with the observed data. Parameterizations were specifically targeted to result in a disseminated infection across all simulations, but not necessarily leading to salivary gland infection in every instance (Figure 4C). From the baseline scenario, the impact of varying parameterizations was demonstrated by running the model 100 times with *G_v_* at 10^6^ PFU/ml. Throughout the scenarios, *μ* at 1/72 h^-1^, *β* at 10^-3^, the midgut growth rate (*r_M_*) at 0.04 h^-1^, the escape rate from the midgut to the hemocoel (*λ_M:H_*) at 0.0005 h^-1^, and *k* of all tissues at 10^20^ PFU/ml, were kept constant. For scenarios two through five, where the value of one parameter was reduced by a tenth to illustrate the impact on model outputs, variations were introduced in the hemocoel growth rate (*r_H_*) and the escape rate from hemocoel to salivary glands (*λ_H:S_*), using Gamma distributions. For *r_H_*, a shape (*α*) of 400 and a scale (*θ*) of 0.025 were used, resulting in an average of 0.04/10 and a variance of 0.00001. For *λ_H:S_*, *α* of 2500 and *θ* of 0.0004 were utilized, achieving an average of 0.0005/10 and a variance of 0.00000002. In scenarios three and five, parameter values were randomly selected from these distributions for each simulation. Additionally, in scenario five, the variance of the Gamma distribution was varied, with simulations run using a mean of 0.00008 and variances of 10^-7.5^, 10^-7.3^, 10^-7^, and 10^-6.5^.

## Author contributions

Conceptualization: S.T., X.M., L.L.

Methodology: J.S.L., M.B.B.

Formal analysis: S.T., J.S.L., M.B.B., E.S.-L., L.L.

Investigation: S.T., M.L., M.P., A.L.

Resources: C.T.D., Oum.F., Ous.F., A.A.S.

Data curation: E.S.-L.

Writing – original draft: S.T., L.L.

Writing – review & editing: S.T., J.S.L., M.B.B., E.S.-L., X.M., L.L.

Visualization: S.T., J.S.L.

Supervision: L.L.

Funding acquisition: S.T., J.S.L., E.S.-L., X.M., L.L.

## Acknowledgments

We thank Catherine Lallemand for assistance with mosquito rearing. We thank Caroline Manet and all members of the Lambrechts lab for their input during the project. We are grateful to Claudia Romero-Vivas and Silvânia Da Veiga Leal, who helped establish the mosquito colonies from Colombia and Cape Verde, respectively. We thank John-Paul Mutebi, Noah Rose, and Lindy McBride for initially providing the colony from Uganda. We thank Rick Jarman for providing the ZIKV strain from Thailand.

## Supplementary materials

**Figure S1. Schematic representation of experimental approaches.**

**(A)** Overview of the circular polymerase extension reaction (CPER) method for creating chimeric ZIKV. Seven fragments spanning the full-length ZIKV genome, including a promoter sequence with HDVr and pA signals, were amplified by PCR. These fragments were assembled using the CPER technique, and the resulting products were transfected into Huh-7 cells. The virus was subsequently propagated through passaging in C6/36 cells. (**B**) Design of the *in vivo* mosquito experiments. *Aedes aegypti* mosquitoes were offered artificial infectious blood meals via a membrane-feeding system. Saliva samples were collected on days 7, 10, and 14 post infectious blood meal to detect infectious virus by focus-forming assay. Additionally, mosquito bodies and heads were dissected to detect viral RNA by RT-PCR. (**C**) Comparative genomic analysis. Full-length viral genome sequences of the iSenegal and iThailand strains were compared to highlight genetic differences at the nucleotide and amino-acid levels.

**Figure S2. Natural and engineered strains display similar *in vivo* phenotypes in mosquitoes.**

Barplots show the infection prevalence (**A**), dissemination prevalence (**B**), and transmission efficiency (**C**) in *Ae. aegypti* mosquitoes from Colombia, following an infectious blood meal containing an expected virus titer of 1×10⁶ PFU/ml. Infection prevalence is the proportion of blood-fed mosquitoes with a virus-positive body (measured by RT-PCR). Dissemination prevalence is the proportion of infected mosquitoes with a virus-positive head (measured by RT-PCR). Transmission efficiency is the proportion of blood-fed mosquitoes with infectious saliva (measured by focus-forming assay). Bars are color-coded to represent the different virus strains, and data are presented as proportions ±95% confidence intervals. The corresponding raw data are provided in Table S3.

**Figure S3. Both structural and non-structural genes influence the probability of mosquito infection by rSenegal and rThailand strains at low infectious doses.**

**(A)** Dose-response curves representing the infection prevalence in *Ae. aegypti* mosquitoes 3 days after oral exposure to varying doses of the first set of chimeric viruses. Infection prevalence is plotted as a function of the oral infectious dose (blood meal titer). (**B**) Oral infectious dose required to infect 50% of mosquitoes (OID₅₀) estimates are shown for the first set of chimeric viruses, along with 95% confidence intervals derived from logistic regression fits. Data points and logistic regression lines are color-coded according to the viruses. The corresponding raw data are provided in Table S3.

**Figure S4. Asymmetric effect of individual genes from rSenegal and rThailand strains on viral growth in mosquito cells.**

(**A**-**B**) Schematic representations of the second (**A**) and third (**B**) sets of chimeric viruses. (**C**-**D**) Plaque diameters were measured on Vero E6 cell monolayers infected with the second (**C**) and third (**D**) sets of chimeric viruses. Measurements were normalized to the respective parental virus strains and quantified using ImageJ software. (**E**-**F**) Viral growth kinetics in C6/36 cells infected with chimeric viruses from the second (**E**) and third (**F**) sets at a MOI of 0.01. Infectious virus titers were determined using a plaque assay on supernatants collected at regular intervals. (**C**-**F**) Data are presented as mean ±SEM from 3-7 biological replicates, with statistical significance determined by one-way ANOVA followed by Dunnett’s test (\**p* <0.05; \*\**p* <0.01; ns: non-significant). Lines and bars are color-coded to represent the different chimeric viruses, and the parental strains are depicted with thicker lines.

**Figure S5. Structural genes influence virus transmission irrespective of mosquito genetic background.**

*Aedes aegypti* mosquitoes from Uganda (**A**) and Cape Verde (**B**) were exposed to the first set of chimeric viruses via an infectious blood meal. Mosquitoes were collected on days 7, 10, and 14 post infectious blood meal to assess the transmission efficiency, defined as the proportion of blood-fed *Ae. aegypti* mosquitoes with infectious saliva (measured by focus-forming assay). Lines and data points are color-coded to represent the different chimeric viruses, and the parental strains are depicted with thicker lines. Each data point reflects the observed proportions, with the size of the point proportional to the number of mosquitoes tested in each group. Logistic regression results are depicted as fitted lines, with error bars indicating 95% confidence intervals for the fits. Raw data and logistic regression outcomes are provided in Table S5.

**Figure S6. Stochastic models of *in vitro* viral growth kinetics and *in vivo* mosquito infection dynamics reproduce the experimental data.**

**(A)** Kinetics of viral growth *in vitro* (expressed in log_10_-transformed PFU/ml), showing the model output overlaid onto the experimental data. The solid lines represent the model results for two combinations of growth rates (*r*, expressed in h^-1^) and starting virus concentrations (*s*, expressed in log_10_-transformed PFU/ml), with a carrying capacity (*k*) fixed at 10^8.5^ PFU/ml. Red circles and blue triangles represent the experimental data for the rSenegal and rThailand strains, respectively. (**B**) Dose-response curves of mosquito infection *in vivo*, showing the model output overlaid onto the experimental data. The dashed lines represent the model results for three values of midgut infection probability (*β*, expressed in log_10_-transformed PFU/ml). Red circles and blue triangles represent the experimental data and their 95% confidence intervals for the rSenegal and rThailand strains, respectively.

**Figure S7. Sensitivity analysis for the stochastic model of midgut infection.**

Output from Latin hypercube sampling, using the following ranges for parameter values: blood meal clearance rate (*μ*) 1/72 – 1/12 h^-1^, probability of midgut infection (*β*) 0 – 0.0001, growth rate (*r*) 0.001 – 0.1 h^-1^, and carrying capacity (*k*) 10^3^ – 10^20^ PFU/ml.

**Figure S8. Sensitivity analysis for the stochastic model of viral dissemination and transmission potential.**

Outputs from Latin hypercube sampling for the hemocoel at seven days (**A**) and the salivary glands at 10 days (**B**) post infectious blood meal, using the following ranges for parameter values: blood meal clearance rate (*μ*) 0.014 – 0.08 h^-1^, probability of midgut infection (*β*) 10^-4.7^ – 10^-4^, growth rate (*r*) 0.001 – 0.1 h^-1^, carrying capacity (*k*) 10^3^ – 10^20^ PFU/ml, and escape rate (*λ*) 10^-8^ – 10^-5^ h^-1^.

**Table S1. Primer pairs used for the construction of chimeric viruses.**

**Table S2. Oligonucleotide sequences.**

**Table S3. Raw data of experimental mosquito infections.**

**Table S4. Statistical analysis of viral infection, dissemination, and transmission in mosquitoes.** The table shows the results of the multivariate logistic regression of the proportion of blood-fed mosquitoes with a midgut infection (infection prevalence), the proportion of midgut-infected mosquitoes with a disseminated infection (dissemination prevalence), and the proportion of mosquitoes with a disseminated infection that expectorated virus in their saliva (transmission prevalence). The colony of mosquitoes and the panel of chimeric viruses are indicated in the two left columns. For each phenotype, the minimal adequate model was obtained by sequentially removing non-significant terms (*p >*0.05) from the full-factorial model. Time was considered a continuous variable and the dose (blood meal titer) was log_10_-transformed. Df: degrees of freedom; LR: likelihood ratio.

**Table S5. Raw data of experimental mosquito infections and statistical analysis of transmission efficiency in mosquitoes**. The table shows the results of the multivariate logistic regression of the proportion of blood-fed mosquitoes that expectorated virus in their saliva (transmission efficiency). The colony of mosquitoes and the panel of chimeric viruses are indicated. The minimal adequate model was obtained by sequentially removing non-significant terms (*p* <0.05) from the full-factorial model. Df: degrees of freedom; LR: likelihood ratio.

