## Supplemental figures for "Polygenic viral factors enable efficient mosquito-borne transmission of African Zika virus"

Sup Fig. 1 Torii et al.

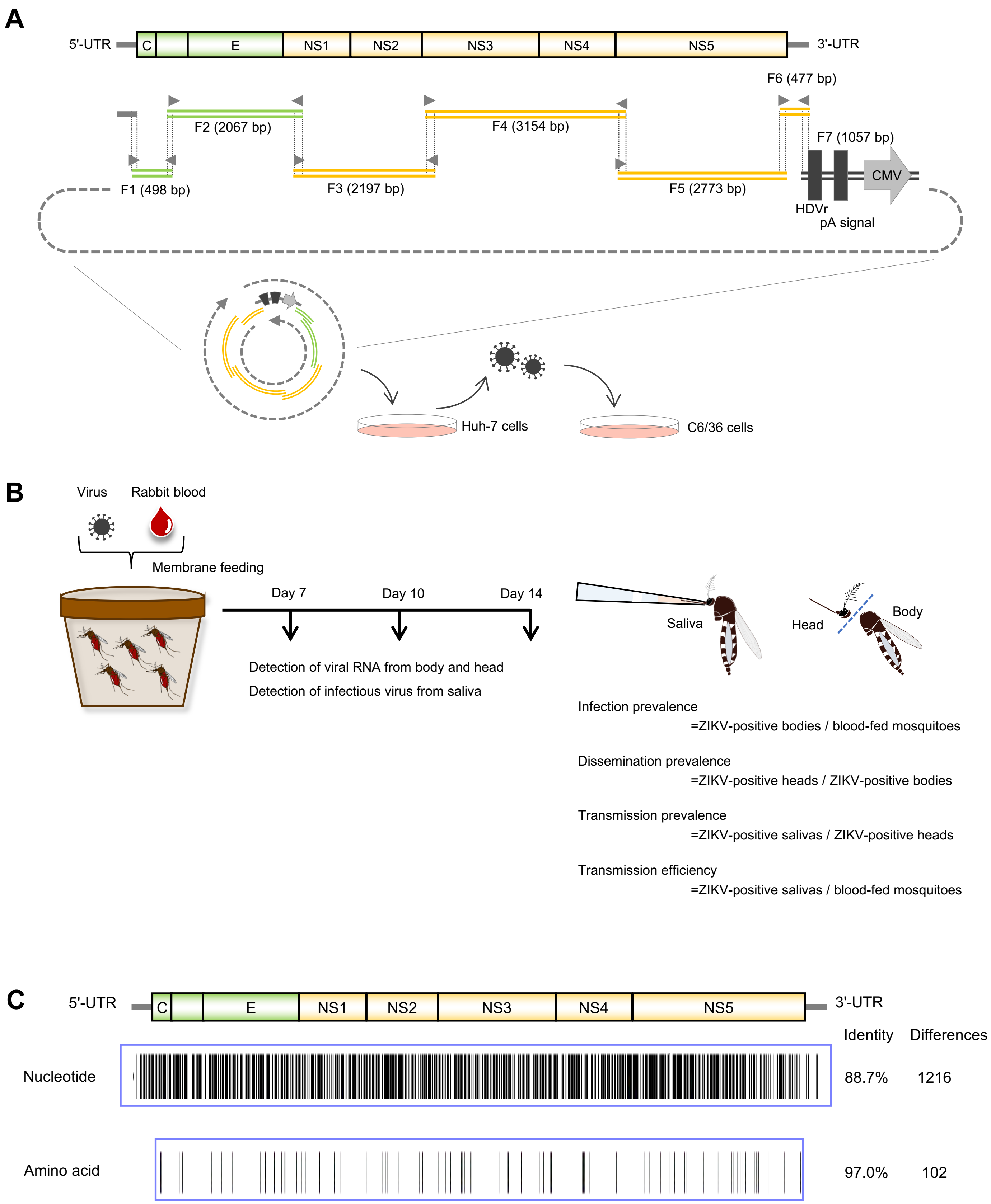

Sup Fig. 2 Torii et al.

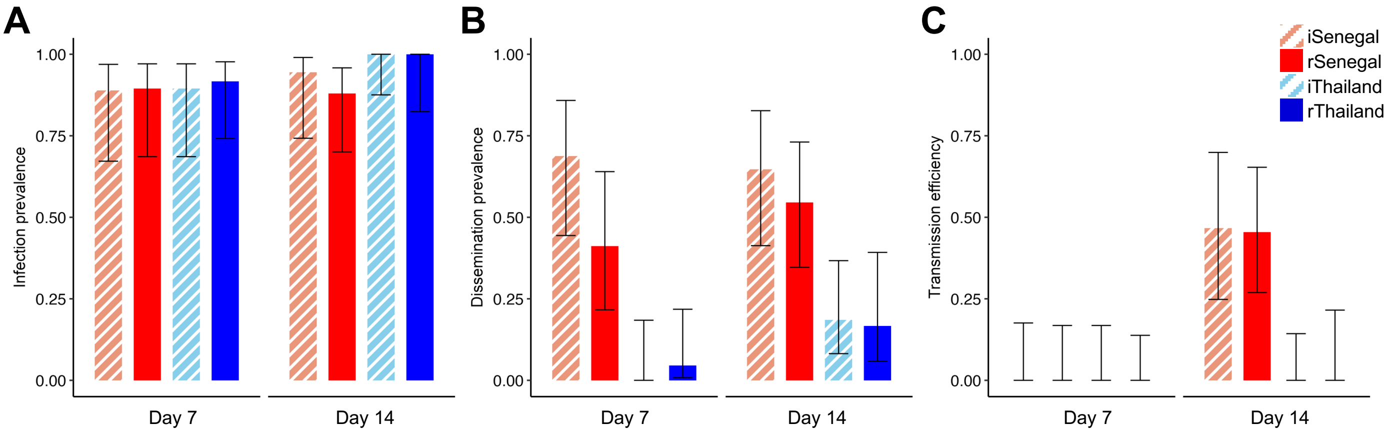

Sup Fig. 3 Torii et al.

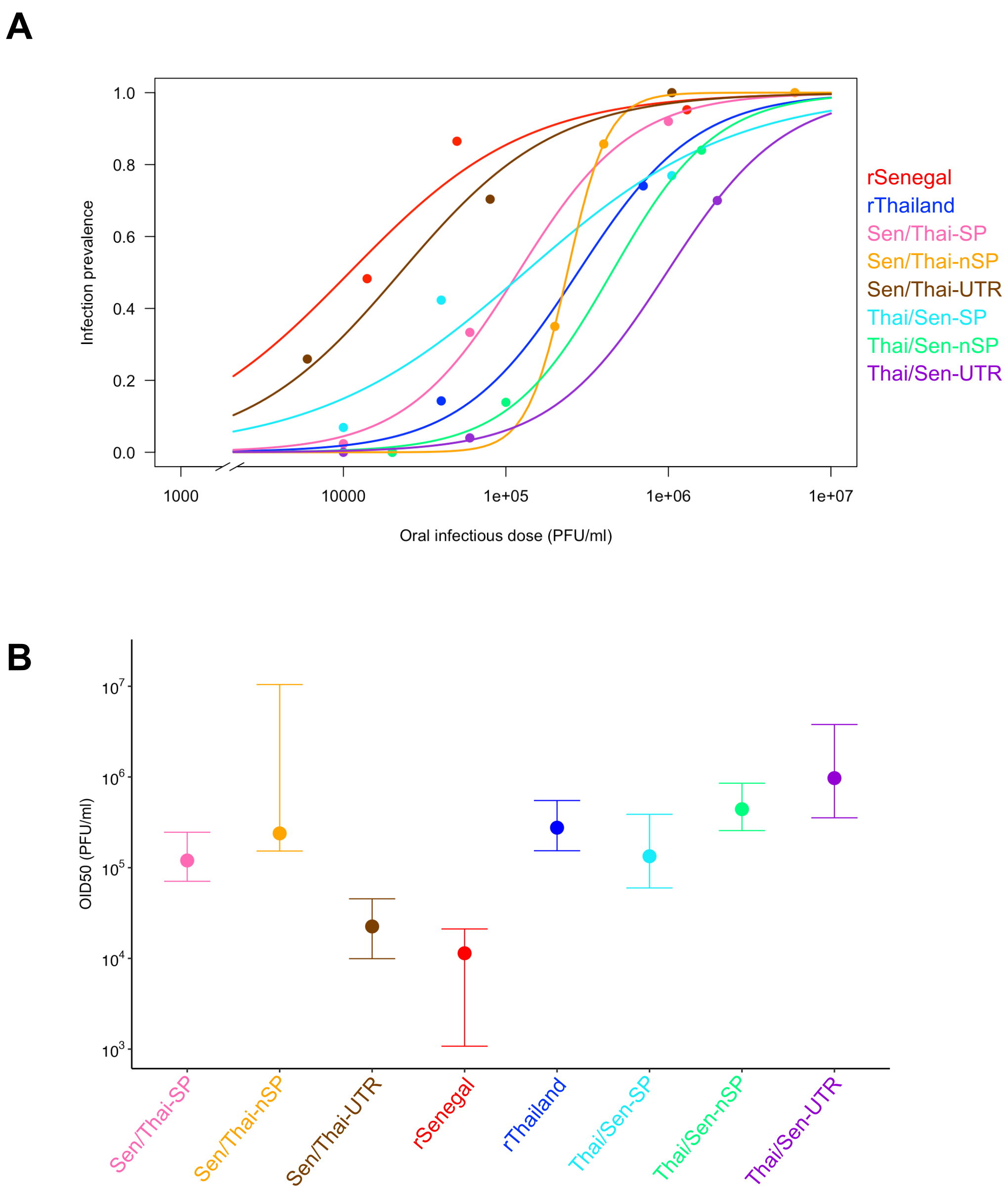

Sup Fig. 4 Torii et al.

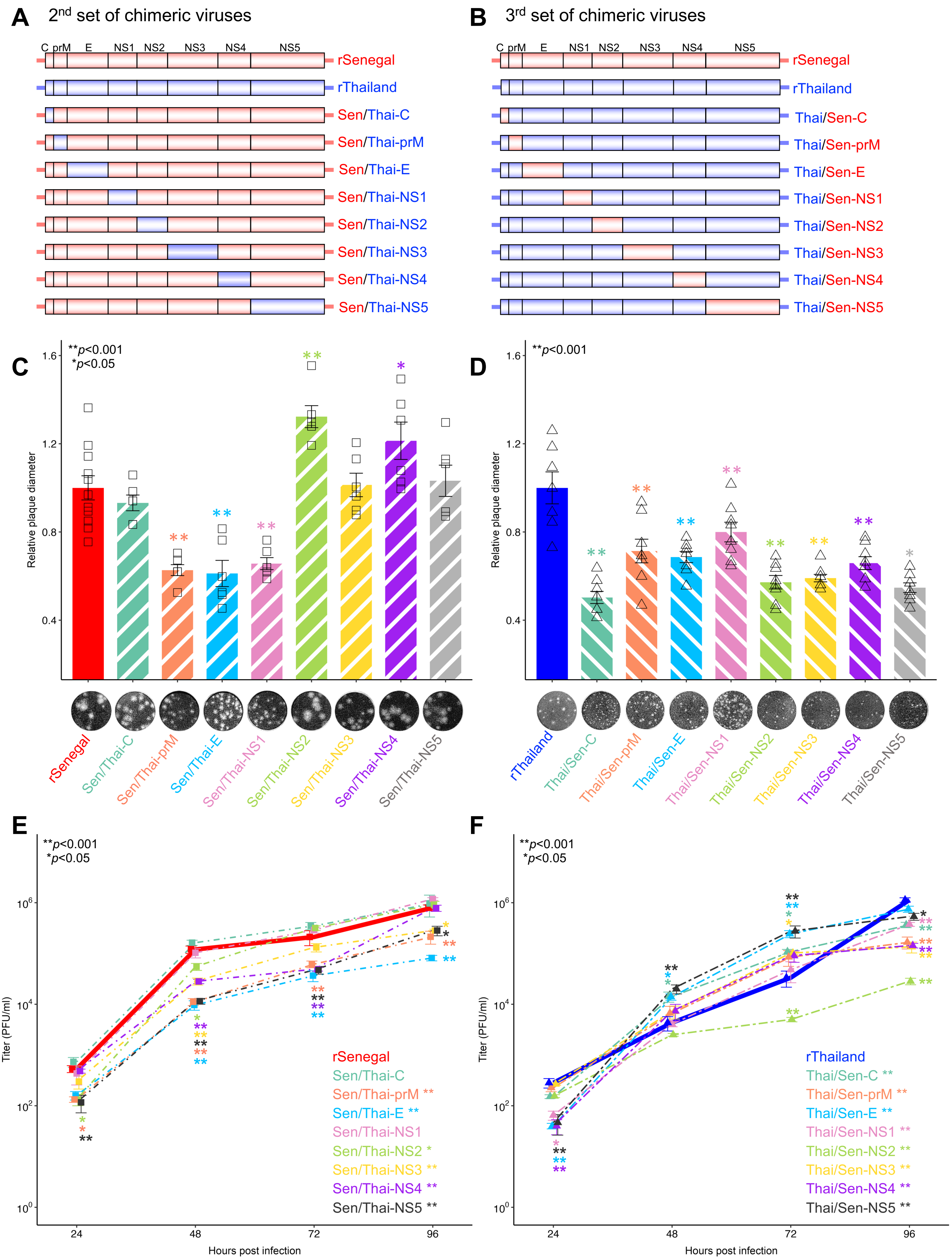

Sup Fig. 5 Torii et al.

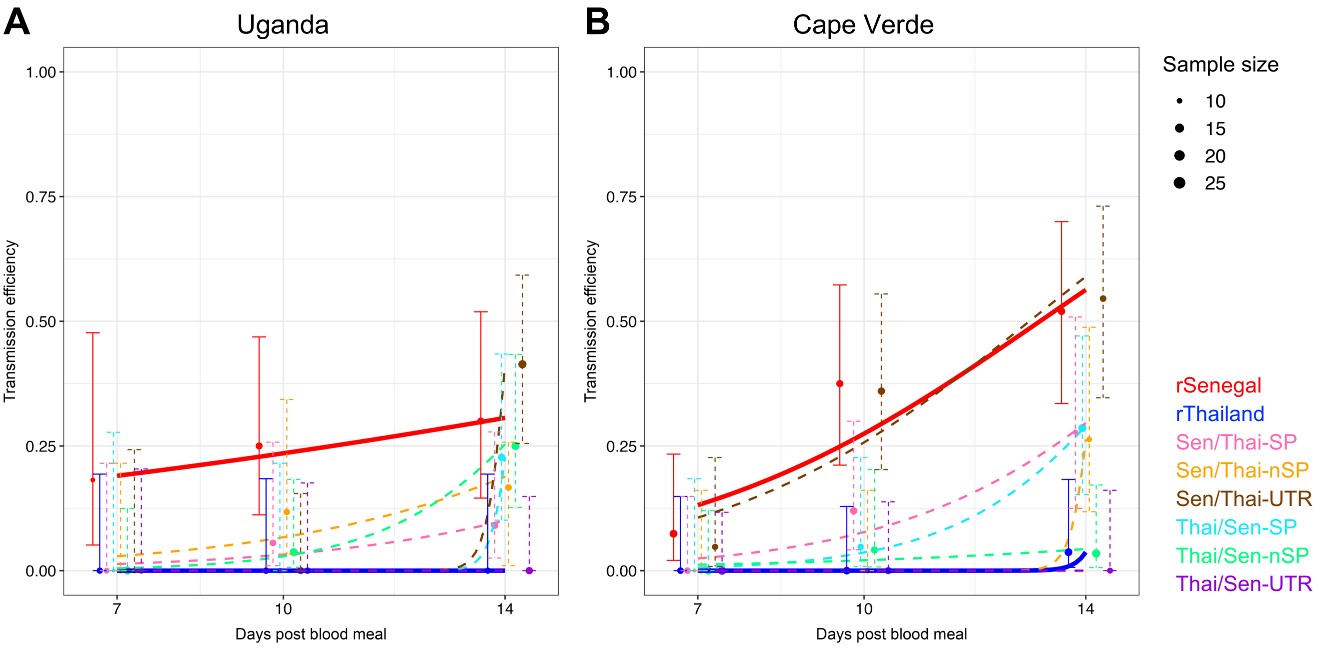

Sup Fig. 6 Torii et al.

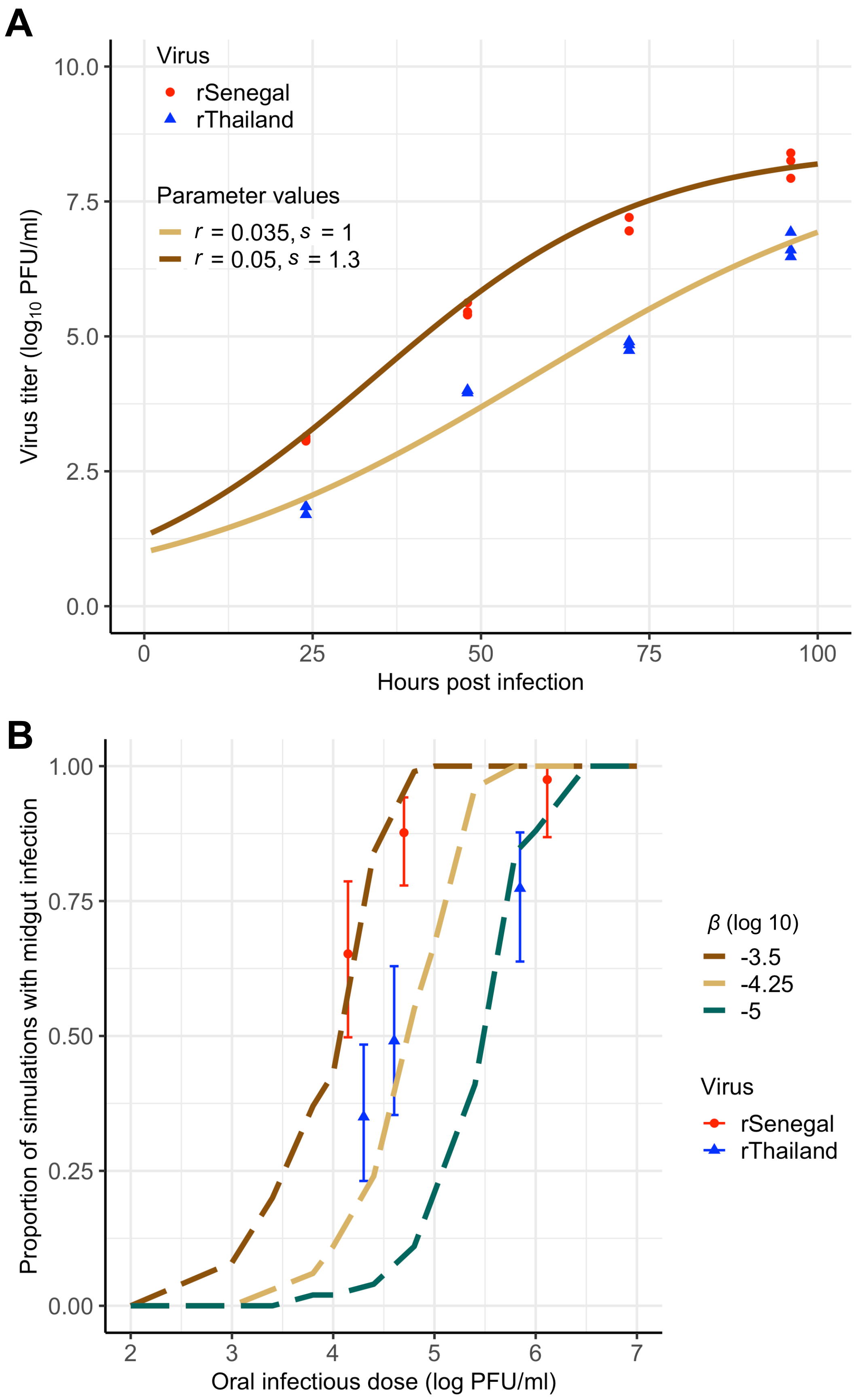

Sup Fig. 7 Torii et al.

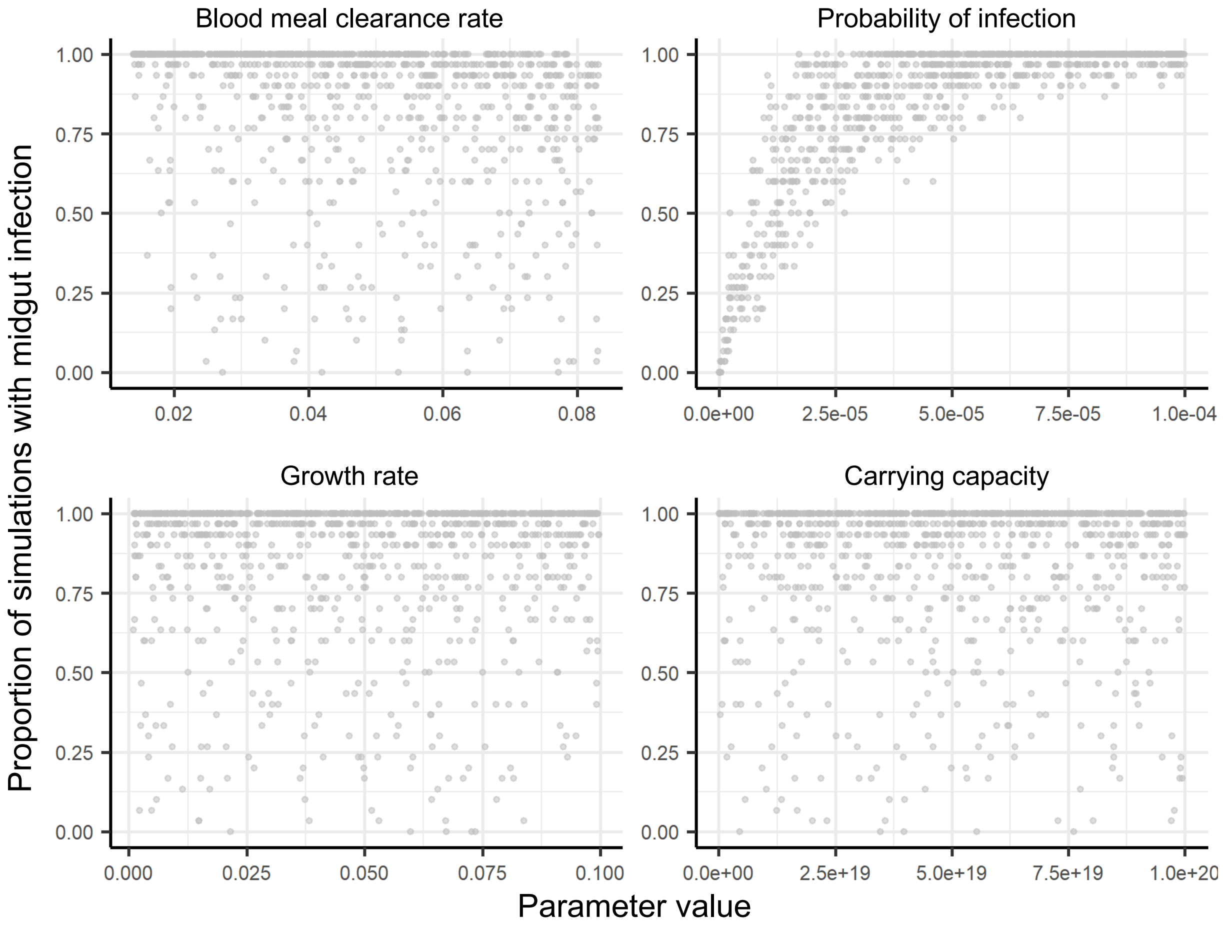

Sup Fig. 8 Torii et al.

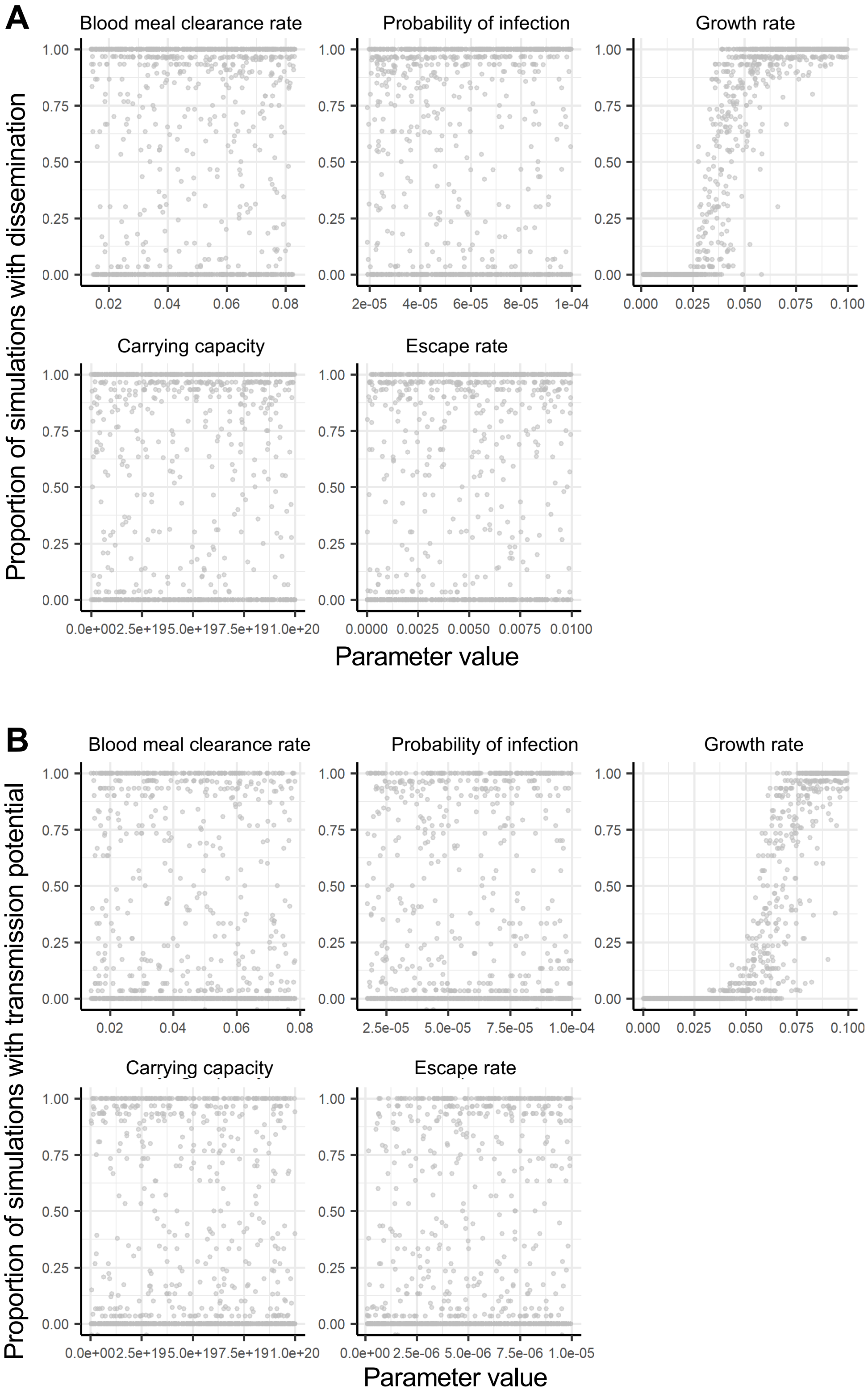
