## Supplemental tables for "Polygenic viral factors enable efficient mosquito-borne transmission of African Zika virus"

Table S1. Primer pairs used for the construction of chimeric viruses.

| Chimeric virus | Fragment | Forward primer | Reverse primer | Templates |
|---|---|---|---|---|
| rSenegal | F1 | Sen-Linker/F1-F | Sen-F1/F2-R | Senegal |
| | F2 | Sen-F1/F2-F | Sen-F2/F3-R | Senegal |
| | F3 | Sen-F2/F3-F | Sen-F3/F4-R | Senegal |
| | F4 | Sen-F3/F4-F | Sen-F4/F5-R | Senegal |
| | F5 | Sen-F4/F5-F | Sen-F5/F6-R | Senegal |
| | F6 | Sen-F5/F6-F | Sen-F6/Linker-R | Senegal |
| | F7 | Sen-F6/Linker-F | Sen-Linker/F1-R | Linker |
| rThailand | F1 | Thai-Linker/F1-F | Thai-F1/F2-R | Thailand |
| | F2 | Thai-F1/F2-F | Thai-F2/F3-R | Thailand |
| | F3 | Thai-F2/F3-F | Thai-F3/F4-R | Thailand |
| | F4 | Thai-F3/F4-F | Thai-F4/F5-R | Thailand |
| | F5 | Thai-F4/F5-F | Thai-F5/F6-R | Thailand |
| | F6 | Thai-F5/F6-F | Thai-F6/Linker-R | Thailand |
| | F7 | Thai-F6/Linker-F | Thai-Linker/F1-R | Linker |
| Sen/Thai-SP | F1-1 | Sen-Linker/F1-F | Sen/Thai-SP-F1-1-R | Senegal |
| | F1-2 | Sen/Thai-SP-F1-1-F | Thai-F1/F2-R | Thailand |
| | F2 | Thai-F1/F2-F | Sen/Thai-SP-F2/F3-R | Thailand |
| | F3 | Sen/Thai-SP-F2/F3-F | Sen-F3/F4-R | Senegal |
| | F4 | Sen-F3/F4-F | Sen-F4/F5-R | Senegal |
| | F5 | Sen-F4/F5-F | Sen-F5/F6-R | Senegal |
| | F6 | Sen-F5/F6-F | Sen-F6/Linker-R | Senegal |
| | F7 | Sen-F6/Linker-F | Sen-Linker/F1-R | Linker |
| Thai/Sen-SP | F1-1 | Thai-Linker/F1-F | Sen/Thai-SP-F1-1-R | Thailand |
| | F1-2 | Sen/Thai-SP-F1-1-F | Sen-F1/F2-R | Senegal |
| | F2 | Sen-F1/F2-F | Thai/Sen-SP-F2/F3-R | Senegal |
| | F3 | Thai/Sen-SP-F2/F3-F | Thai-F3/F4-R | Thailand |
| | F4 | Thai-F3/F4-F | Thai-F4/F5-R | Thailand |
| | F5 | Thai-F4/F5-F | Thai-F5/F6-R | Thailand |
| | F6 | Thai-F5/F6-F | Thai-F6/Linker-R | Thailand |
| | F7 | Thai-F6/Linker-F | Thai-Linker/F1-R | Linker |
| Sen/Thai-nSP | F1 | Sen-Linker/F1-F | Sen-F1/F2-R | Senegal |
| | F2 | Sen-F1/F2-F | Thai/Sen-SP-F2/F3-R | Senegal |
| | F3 | Thai/Sen-SP-F2/F3-F | Thai-F3/F4-R | Thailand |
| | F4 | Thai-F3/F4-F | Thai-F4/F5-R | Thailand |
| | F5 | Thai-F4/F5-F | Sen/Thai-nSP-F5/F6-R | Thailand |
| | F6 | Sen/Thai-nSP-F5/F6-F | Sen-F6/Linker-R | Senegal |
| | F7 | Sen-F6/Linker-F | Sen-Linker/F1-R | Linker |
| Thai/Sen-nSP | F1 | Thai-Linker/F1-F | Thai-F1/F2-R | Thailand |
| | F2 | Thai-F1/F2-F | Sen/Thai-SP-F2/F3-R | Thailand |
| | F3 | Sen/Thai-SP-F2/F3-F | Sen-F3/F4-R | Senegal |
| | F4 | Sen-F3/F4-F | Sen-F4/F5-R | Senegal |
| | F5 | Sen-F4/F5-F | Thai/Sen-nSP-F5/F6-R | Senegal |
| | F6 | Thai/Sen-nSP-F5/F6-F | Thai-F6/Linker-R | Thailand |
| | F7 | Thai-F6/Linker-F | Thai-Linker/F1-R | Linker |
| Sen/Thai-UTR | F1-1 | Thai-Linker/F1-F | Sen/Thai-SP-F1-1-R | Thailand |
| | F1-2 | Sen/Thai-SP-F1-1-F | Sen-F1/F2-R | Senegal |
| | F2 | Sen-F1/F2-F | Sen-F2/F3-R | Senegal |
| | F3 | Sen-F2/F3-F | Sen-F3/F4-R | Senegal |
| | F4 | Sen-F3/F4-F | Sen-F4/F5-R | Senegal |
| | F5 | Sen-F4/F5-F | Thai/Sen-nSP-F5/F6-R | Senegal |
| | F6 | Thai/Sen-nSP-F5/F6-F | Thai-F6/Linker-R | Thailand |
| | F7 | Thai-F6/Linker-F | Thai-Linker/F1-R | Linker |
| Thai/Sen-UTR | F1-1 | Sen-Linker/F1-F | Sen/Thai-SP-F1-1-R | Senegal |
| | F1-2 | Sen/Thai-SP-F1-1-F | Thai-F1/F2-R | Thailand |
| | F2 | Thai-F1/F2-F | Thai-F2/F3-R | Thailand |

| | | | | |
|---|---|---|---|---|
| | F3 | Thai-F2/F3-F | Thai-F3/F4-R | Thailand |
| | F4 | Thai-F3/F4-F | Thai-F4/F5-R | Thailand |
| | F5 | Thai-F4/F5-F | Sen/Thai-nSP-F5/F6-R | Thailand |
| | F6 | Sen/Thai-nSP-F5/F6-F | Sen-F6/Linker-R | Senegal |
| | F7 | Sen-F6/Linker-F | Sen-Linker/F1-R | Linker |
| Sen/Thai-C | F1-1 | Sen-Linker/F1-F | Sen/Thai-SP-F1-1-R | Senegal |
| | F1-2 | Sen/Thai-SP-F1-1-F | Sen/Thai-C-F1/F2-R | Thailand |
| | F2 | Sen/Thai-C-F1/F2-F | Sen-F2/F3-R | Senegal |
| | F3 | Sen-F2/F3-F | Sen-F3/F4-R | Senegal |
| | F4 | Sen-F3/F4-F | Sen-F4/F5-R | Senegal |
| | F5 | Sen-F4/F5-F | Sen-F5/F6-R | Senegal |
| | F6 | Sen-F5/F6-F | Sen-F6/Linker-R | Senegal |
| | F7 | Sen-F6/Linker-F | Sen-Linker/F1-R | Linker |
| Sen/Thai-prM | F1 | Sen-Linker/F1-F | Sen/Thai-prM-F1/F2-R | Senegal |
| | F2-1 | Sen/Thai-prM-F1/F2-F | Sen/Thai-prM-F2-1-R | Thailand |
| | F2-2 | Sen/Thai-prM-F2-1-F | Sen-F2/F3-R | Senegal |
| | F3 | Sen-F2/F3-F | Sen-F3/F4-R | Senegal |
| | F4 | Sen-F3/F4-F | Sen-F4/F5-R | Senegal |
| | F5 | Sen-F4/F5-F | Sen-F5/F6-R | Senegal |
| | F6 | Sen-F5/F6-F | Sen-F6/Linker-R | Senegal |
| | F7 | Sen-F6/Linker-F | Sen-Linker/F1-R | Linker |
| Sen/Thai-E | F1 | Sen-Linker/F1-F | Sen-F1/F2-R | Senegal |
| | F2-1 | Sen-F1/F2-F | Sen/Thai-E-F2-1-R | Senegal |
| | F2-2 | Sen/Thai-E-F2-1-F | Sen/Thai-SP-F2/F3-R | Thailand |
| | F3 | Sen/Thai-SP-F2/F3-F | Sen-F3/F4-R | Senegal |
| | F4 | Sen-F3/F4-F | Sen-F4/F5-R | Senegal |
| | F5 | Sen-F4/F5-F | Sen-F5/F6-R | Senegal |
| | F6 | Sen-F5/F6-F | Sen-F6/Linker-R | Senegal |
| | F7 | Sen-F6/Linker-F | Sen-Linker/F1-R | Linker |
| Sen/Thai-NS1 | F1 | Sen-Linker/F1-F | Sen-F1/F2-R | Senegal |
| | F2 | Sen-F1/F2-F | Thai/Sen-SP-F2/F3-R | Senegal |
| | F3-1 | Thai/Sen-SP-F2/F3-F | Sen/Thai-NS1-F3-1-R | Thailand |
| | F3-2 | Sen/Thai-NS1-F3-1-F | Sen-F3/F4-R | Senegal |
| | F4 | Sen-F3/F4-F | Sen-F4/F5-R | Senegal |
| | F5 | Sen-F4/F5-F | Sen-F5/F6-R | Senegal |
| | F6 | Sen-F5/F6-F | Sen-F6/Linker-R | Senegal |
| | F7 | Sen-F6/Linker-F | Sen-Linker/F1-R | Linker |
| Sen/Thai-NS2 | F1 | Sen-Linker/F1-F | Sen-F1/F2-R | Senegal |
| | F2 | Sen-F1/F2-F | Sen-F2/F3-R | Senegal |
| | F3-1 | Sen-F2/F3-F | Sen/Thai-NS2-F3-1-R | Senegal |
| | F3-2 | Sen/Thai-NS2-F3-1-F | Sen/Thai-NS2-F3/F4-R | Thailand |
| | F4 | Sen/Thai-NS2-F3/F4-F | Sen-F4/F5-R | Senegal |
| | F5 | Sen-F4/F5-F | Sen-F5/F6-R | Senegal |
| | F6 | Sen-F5/F6-F | Sen-F6/Linker-R | Senegal |
| | F7 | Sen-F6/Linker-F | Sen-Linker/F1-R | Linker |
| Sen/Thai-NS3 | F1 | Sen-Linker/F1-F | Sen-F1/F2-R | Senegal |
| | F2 | Sen-F1/F2-F | Sen-F2/F3-R | Senegal |
| | F3 | Sen-F2/F3-F | Sen/Thai-NS3-F3/F4-R | Senegal |
| | F4-1 | Sen/Thai-NS3-F3/F4-F | Sen/Thai-NS3-F4-1-R | Thailand |
| | F4-2 | Sen/Thai-NS3-F4-1-F | Sen-F4/F5-R | Senegal |
| | F5 | Sen-F4/F5-F | Sen-F5/F6-R | Senegal |
| | F6 | Sen-F5/F6-F | Sen-F6/Linker-R | Senegal |
| | F7 | Sen-F6/Linker-F | Sen-Linker/F1-R | Linker |
| Sen/Thai-NS4 | F1 | Sen-Linker/F1-F | Sen-F1/F2-R | Senegal |
| | F2 | Sen-F1/F2-F | Sen-F2/F3-R | Senegal |
| | F3 | Sen-F2/F3-F | Sen-F3/F4-R | Senegal |
| | F4-1 | Sen-F3/F4-F | Sen/Thai-NS4-F4-1-R | Senegal |
| | F4-2 | Sen/Thai-NS4-F4-1-F | Sen/Thai-NS4-F4/F5-R | Thailand |

| | | | | |
|---|---|---|---|---|
| | F5 | Sen/Thai-NS4-F4/F5-F | Sen-F5/F6-R | Senegal |
| | F6 | Sen-F5/F6-F | Sen-F6/Linker-R | Senegal |
| | F7 | Sen-F6/Linker-F | Sen-Linker/F1-R | Linker |
| Sen/Thai-NS5 | F1 | Sen-Linker/F1-F | Sen-F1/F2-R | Senegal |
| | F2 | Sen-F1/F2-F | Sen-F2/F3-R | Senegal |
| | F3 | Sen-F2/F3-F | Sen-F3/F4-R | Senegal |
| | F4 | Sen-F3/F4-F | Sen/Thai-NS5-F4/F5-R | Senegal |
| | F5 | Sen/Thai-NS5-F4/F5-F | Sen/Thai-nSP-F5/F6-R | Thailand |
| | F6 | Sen/Thai-nSP-F5/F6-F | Sen-F6/Linker-R | Merge |
| | F7 | Sen-F6/Linker-F | Sen-Linker/F1-R | Linker |
| Thai/Sen-C | F1-1 | Thai-Linker/F1-F | Sen/Thai-SP-F1-1-R | Thailand |
| | F1-2 | Sen/Thai-SP-F1-1-F | Sen/Thai-prM-F1/F2-R | Senegal |
| | F2 | Sen/Thai-prM-F1/F2-F | Thai-F2/F3-R | Thailand |
| | F3 | Thai-F2/F3-F | Thai-F3/F4-R | Thailand |
| | F4 | Thai-F3/F4-F | Thai-F4/F5-R | Thailand |
| | F5 | Thai-F4/F5-F | Thai-F5/F6-R | Thailand |
| | F6 | Thai-F5/F6-F | Thai-F6/Linker-R | Thailand |
| | F7 | Thai-F6/Linker-F | Thai-Linker/F1-R | Linker |
| Thai/Sen-prM | F1 | Thai-Linker/F1-F | Sen/Thai-C-F1/F2-R | Thailand |
| | F2-1 | Sen/Thai-C-F1/F2-F | Sen/Thai-E-F2-1-R | Senegal |
| | F2-2 | Sen/Thai-E-F2-1-F | Thai-F2/F3-R | Thailand |
| | F3 | Thai-F2/F3-F | Thai-F3/F4-R | Thailand |
| | F4 | Thai-F3/F4-F | Thai-F4/F5-R | Thailand |
| | F5 | Thai-F4/F5-F | Thai-F5/F6-R | Thailand |
| | F6 | Thai-F5/F6-F | Thai-F6/Linker-R | Thailand |
| | F7 | Thai-F6/Linker-F | Thai-Linker/F1-R | Linker |
| Thai/Sen-E | F1 | Thai-Linker/F1-F | Thai-F1/F2-R | Thailand |
| | F2-1 | Thai-F1/F2-F | Sen/Thai-prM-F2-1-R | Thailand |
| | F2-2 | Sen/Thai-prM-F2-1-F | Thai/Sen-SP-F2/F3-R | Senegal |
| | F3 | Thai/Sen-SP-F2/F3-F | Thai-F3/F4-R | Thailand |
| | F4 | Thai-F3/F4-F | Thai-F4/F5-R | Thailand |
| | F5 | Thai-F4/F5-F | Thai-F5/F6-R | Thailand |
| | F6 | Thai-F5/F6-F | Thai-F6/Linker-R | Thailand |
| | F7 | Thai-F6/Linker-F | Thai-Linker/F1-R | Linker |
| Thai/Sen-NS1 | F1 | Thai-Linker/F1-F | Thai-F1/F2-R | Thailand |
| | F2 | Thai-F1/F2-F | Sen/Thai-SP-F2/F3-R | Thailand |
| | F3-1 | Sen/Thai-SP-F2/F3-F | Sen/Thai-NS2-F3-1-R | Senegal |
| | F3-2 | Sen/Thai-NS2-F3-1-F | Thai-F3/F4-R | Thailand |
| | F4 | Thai-F3/F4-F | Thai-F4/F5-R | Thailand |
| | F5 | Thai-F4/F5-F | Thai-F5/F6-R | Thailand |
| | F6 | Thai-F5/F6-F | Thai-F6/Linker-R | Thailand |
| | F7 | Thai-F6/Linker-F | Thai-Linker/F1-R | Linker |
| Thai/Sen-NS2 | F1 | Thai-Linker/F1-F | Thai-F1/F2-R | Thailand |
| | F2 | Thai-F1/F2-F | Thai-F2/F3-R | Thailand |
| | F3-1 | Thai-F2/F3-F | Sen/Thai-NS1-F3-1-R | Thailand |
| | F3-2 | Sen/Thai-NS1-F3-1-F | Sen/Thai-NS3-F3/F4-R | Senegal |
| | F4 | Sen/Thai-NS3-F3/F4-F | Thai-F4/F5-R | Thailand |
| | F5 | Thai-F4/F5-F | Thai-F5/F6-R | Thailand |
| | F6 | Thai-F5/F6-F | Thai-F6/Linker-R | Thailand |
| | F7 | Thai-F6/Linker-F | Thai-Linker/F1-R | Linker |
| Thai/Sen-NS3 | F1 | Thai-Linker/F1-F | Thai-F1/F2-R | Thailand |
| | F2 | Thai-F1/F2-F | Thai-F2/F3-R | Thailand |
| | F3 | Thai-F2/F3-F | Sen/Thai-NS2-F3/F4-R | Thailand |
| | F4-1 | Sen/Thai-NS2-F3/F4-F | Sen/Thai-NS4-F4-1-R | Senegal |
| | F4-2 | Sen/Thai-NS4-F4-1-F | Thai-F4/F5-R | Thailand |
| | F5 | Thai-F4/F5-F | Thai-F5/F6-R | Thailand |
| | F6 | Thai-F5/F6-F | Thai-F6/Linker-R | Thailand |
| | F7 | Thai-F6/Linker-F | Thai-Linker/F1-R | Linker |

| | | | | |
|---|---|---|---|---|
| Thai/Sen-NS4 | F1 | Thai-Linker/F1-F | Thai-F1/F2-R | Thailand |
| | F2 | Thai-F1/F2-F | Thai-F2/F3-R | Thailand |
| | F3 | Thai-F2/F3-F | Thai-F3/F4-R | Thailand |
| | F4-1 | Thai-F3/F4-F | Sen/Thai-NS3-F4-1-R | Thailand |
| | F4-2 | Sen/Thai-NS3-F4-1-F | Sen/Thai-NS5-F4/F5-R | Senegal |
| | F5 | Sen/Thai-NS5-F4/F5-F | Thai-F5/F6-R | Thailand |
| | F6 | Thai-F5/F6-F | Thai-F6/Linker-R | Thailand |
| | F7 | Thai-F6/Linker-F | Thai-Linker/F1-R | Linker |
| Thai/Sen-NS5 | F1 | Thai-Linker/F1-F | Thai-F1/F2-R | Thailand |
| | F2 | Thai-F1/F2-F | Thai-F2/F3-R | Thailand |
| | F3 | Thai-F2/F3-F | Thai-F3/F4-R | Thailand |
| | F4 | Thai-F3/F4-F | Sen/Thai-NS4-F4/F5-R | Thailand |
| | F5 | Sen/Thai-NS4-F4/F5-F | Thai/Sen-nSP-F5/F6-R | Senegal |
| | F6 | Thai/Sen-nSP-F5/F6-F | Thai-F6/Linker-R | Merge |
| | F7 | Thai-F6/Linker-F | Thai-Linker/F1-R | Linker |

Table S2. Oligonuclotide sequences.

| Targets | Reaction | Oligonuclotide name | Sequence (5'-3') |
|---|---|---|---|
| **PCR primers** | | | |
| Zika virus | PCR | ZIKV-PCR-F | GTATGGAATGGAGATAAGGCCC |
| | | ZIKV-PCR-R | ACCAGCACTGCCATTGATGTGC |
| | | ZIKV-PCR-R1 | TCGTATTGCCAACCAGGCCAAAGC |
| **Primers for CPER** | | | |
| Zika virus | PCR | Sen-F1/F2-F | CCTCCTGCTGACCACAGCCATGGCGGCCGAGATCACTAGACGTGGGAGTTC |
| | | Sen-F1/F2-R | GAACTCCCACGTCTAGTGATCTCGGCCGCCATGGCTGTGGTCAGCAGGAGG |
| | | Sen-F2/F3-F | GATTTTCCTTTCCACGGCTGTTTCTGCTGATGTTGGGTGCTCGGTGGACTTCTC |
| | | Sen-F2/F3-R | GAGAAGTCCACCGAGCACCCAACATCAGCAGAAACAGCCGTGGAAAGGAAAATC |
| | | Sen-F3/F4-F | TACCCTTCGCTGCAGGAGCGTGG |
| | | Sen-F3/F4-R | TTTTTCACTTCTTTGGGAGCAGGC |
| | | Sen-F4/F5-F | ACAGTAACAAGAAATGCCGGTCTGG |
| | | Sen-F4/F5-R | TTCAGGCGGGCTTTCCACTTCTC |
| | | Sen-F5/F6-F | GGGTCCACACCTGGAGTGCTGTAAGCACCAATTTCAATGTTGTCAGGCC |
| | | Sen-F5/F6-R | GGCCTGACAACATTGAAATTGGTGCTTACAGCACTCCAGGTGTGGACCC |
| | | Sen-F6/Linker-F | CCAGTGTGGGGAAATCCATGGGTCTGGGTCGGCATGGCATCTCCACCTCC |
| | | Sen-F6/Linker-R | GGAGGTGGAGATGCCATGCCGACCCAGACCCATGGATTTCCCCACACTGG |
| | | Sen-Linker/F1-F | CTATATAAGCAGAGCTCGTTTAGTGAACCGAGTTGTTGATCTGTGTGAATCAGAC |
| | | Sen-Linker/F1-R | GTCTGATTCACACAGATCAACAACTCGGTTCACTAAACGAGCTCTGCTTATATAG |
| | | Thai-F1/F2-F | CCTCCTGCTGACCACAGCTATGGCAGCGGAGGTCACTAGACGTGGGAG |
| | | Thai-F1/F2-R | CTCCCACGTCTAGTGACCTCCGCTGCCATAGCTGTGGTCAGCAGGAGG |
| | | Thai-F2/F3-F | CTTCTTATCCACAGCCGTCTCCGCTGATGTGGGGTGCTCGGTGGACTTC |
| | | Thai-F2/F3-R | GAAGTCCACCGAGCACCCCACATCAGCGGAGACGGCTGTGGATAAGAAG |
| | | Thai-F3/F4-F | CCCTTTGCAGCTGGAGCGTGGTACG |
| | | Thai-F3/F4-R | CCTTTTTTACTTCCTTGGGAGCAGGC |
| | | Thai-F4/F5-F | AATCTACACAGTAACAAGAAACGCTGGC |
| | | Thai-F4/F5-R | TTCAAGCGGGCCTTCCATTTCTC |
| | | Thai-F5/F6-F | GGGTCTACACCTGGAGTGCTATAAGCACCAATCTTAGTGTTGTCAGGCC |
| | | Thai-F5/F6-R | GGCCTGACAACACTAAGATTGGTGCTTATAGCACTCCAGGTGTAGACCC |
| | | Thai-F6/Linker-F | GGTGTGGGGAAATCCATGGGTCTGGGTCGGCATGGCATCTCCACCTCC |
| | | Thai-F6/Linker-R | GGAGGTGGAGATGCCATGCCGACCCAGACCCATGGATTTCCCCACACC |
| | | Thai-Linker/F1-F | CTATATAAGCAGAGCTCGTTTAGTGAACCGAGTTGTTGATCTGTGTGAATCAGACTGC |
| | | Thai-Linker/F1-R | GCAGTCTGATTCACACAGATCAACAACTCGGTTCACTAAACGAGCTCTGCTTATATAG |
| | | Sen/Thai-SP-F1-1-F | GGAAACGAGAGTTTCTGGTCATGAAAAACCC |
| | | Sen/Thai-SP-F1-1-R | GGGTTTTTCATGACCAGAAACTCTCGTTTCC |

| | | | |
|---|---|---|---|
| | | Sen/Thai-SP-F2/F3-F | CTTCTTATCCACAGCCGTCTCCGCTGATGTTGGGTGCTCGGTGGACTTCTC |
| | | Sen/Thai-SP-F2/F3-R | GAGAAGTCCACCGAGCACCCAACATCAGCGGAGACGGCTGTGGATAAGAAG |
| | | Thai/Sen-SP-F2/F3-F | GATTTTCCTTTCCACGGCTGTTTCTGCTGATGTGGGGTGCTCGGTGGACTTC |
| | | Thai/Sen-SP-F2/F3-R | GAAGTCCACCGAGCACCCCACATCAGCAGAAACAGCCGTGGAAAGGAAAATC |
| | | Sen/Thai-nSP-F5/F6-F | GGGTCTACACCTGGAGTGCTATAAGCACCAATTTCAATGTTGTCAGGCC |
| | | Sen/Thai-nSP-F5/F6-R | GGCCTGACAACATTGAAATTGGTGCTTATAGCACTCCAGGTGTAGACCC |
| | | Thai/Sen-nSP-F5/F6-F | GGGTCCACACCTGGAGTGCTGTAAGCACCAATCTTAGTGTTGTCAGGCC |
| | | Thai/Sen-nSP-F5/F6-R | GGCCTGACAACACTAAGATTGGTGCTTACAGCACTCCAGGTGTGGACCC |
| | | Sen/Thai-C-F1/F2-F | CCTCCTGCTGACCACAGCTATGGCAGCCGAGATCACTAGACGTGGGAGTTC |
| | | Sen/Thai-C-F1/F2-R | GAACTCCCACGTCTAGTGATCTCGGCTGCCATAGCTGTGGTCAGCAGGAGG |
| | | Sen/Thai-prM-F1/F2-F | CCTCCTGCTGACCACAGCCATGGCGGCGGAGGTCACTAGACGTGGGAG |
| | | Sen/Thai-prM-F1/F2-R | CTCCCACGTCTAGTGACCTCCGCCGCCATGGCTGTGGTCAGCAGGAGG |
| | | Sen/Thai-prM-F2-1-F | GATACTGCTGATTGCCCCGGCATACAGCATCAGGTGCATAGGAGTTAGCAATAG |
| | | Sen/Thai-prM-F2-1-R | CTATTGCTAACTCCTATGCACCTGATGCTGTATGCCGGGGCAATCAGCAGTATC |
| | | Sen/Thai-E-F2-1-F | GATATTGTTGATTGCCCCGGCATACAGCATCAGGTGCATAGGAGTCAGTAATAGGG |
| | | Sen/Thai-E-F2-1-R | CCCTATTACTGACTCCTATGCACCTGATGCTGTATGCCGGGGCAATCAACAATATC |
| | | Sen/Thai-NS1-F3-1-F | CTTAGTAAGGTCAATGGTGACTGCAGGATCAACCGATCATATGGATCACTTC |
| | | Sen/Thai-NS1-F3-1-R | GAAGTGATCCATATGATCGGTTGATCCTGCAGTCACCATTGACCTTACTAAG |
| | | Sen/Thai-NS2-F3-1-F | CTTAGTAAGGTCTATGGTGACAGCAGGATCAACTGATCACATGGATCACTTTTC |
| | | Sen/Thai-NS2-F3-1-R | GAAAAGTGATCCATGTGATCAGTTGATCCTGCTGTCACCATAGACCTTACTAAG |
| | | Sen/Thai-NS2-F3/F4-F | GGTACGTATACGTGAAAACTGGAAAAAGGAGTGGTGCCCTCTGGGACGTGC |
| | | Sen/Thai-NS2-F3/F4-R | GCACGTCCCAGAGGGCACCACTCCTTTTTCCAGTTTTCACGTATACGTACC |
| | | Sen/Thai-NS3-F3/F4-F | GGTATGTGTATGTAAAGACTGGGAAAAGGAGTGGTGCTCTATGGGATGTGCC |
| | | Sen/Thai-NS3-F3/F4-R | GGCACATCCCATAGAGCACCACTCCTTTTCCCAGTCTTTACATACACATACC |
| | | Sen/Thai-NS3-F4-1-F | CAAGGAGTTTGCCGCTGGGAAAAGAGGAGCGGCTTTGGGAGTAATGG |
| | | Sen/Thai-NS3-F4-1-R | CCATTACTCCCAAAGCCGCTCCTCTTTTCCCAGCGGCAAACTCCTTG |
| | | Sen/Thai-NS4-F4-1-F | CAAAGAATTTGCCGCTGGGAAGAGAGGAGCGGCTTTTGGAGTGATGG |
| | | Sen/Thai-NS4-F4-1-R | CCATCACTCCAAAAGCCGCTCCTCTCTTCCCAGCGGCAAATTCTTTG |
| | | Sen/Thai-NS4-F4/F5-F | CGCTGGCTTGGTCAAGAGACGTGGAGGTGGAACGGGAGAGACCC |
| | | Sen/Thai-NS4-F4/F5-R | GGGTCTCTCCCGTTCCACCTCCACGTCTCTTGACCAAGCCAGCG |
| | | Sen/Thai-NS5-F4/F5-F | GAAATGCCGGTCTGGTTAAGAGACGTGGGGGTGGAACAGGAGAGACCC |
| | | Sen/Thai-NS5-F4/F5-R | GGGTCTCTCCTGTTCCACCCCCACGTCTCTTAACCAGACCGGCATTTC |
| | | Sen/Thai-NS2-F3/F4-R | GCACGTCCCAGAGGGCACCACTCCTTTTTCCAGTTTTCACGTATACGTACC |
| | | Sen/Thai-NS3-F3/F4-F | GGTATGTGTATGTAAAGACTGGGAAAAGGAGTGGTGCTCTATGGGATGTGCC |
| | | Sen/Thai-NS3-F3/F4-R | GGCACATCCCATAGAGCACCACTCCTTTTCCCAGTCTTTACATACACATACC |
| | | Sen/Thai-NS3-F4-1-F | CAAGGAGTTTGCCGCTGGGAAAAGAGGAGCGGCTTTGGGAGTAATGG |
| **RT-qPCR for ZIKV** | | | |

| | | | |
|---|---|---|---|
| African lineage | RT-qPCR | qPCR-Af-F | GTCGCTGTCCAACACAAG |
| | | qPCR-Af-R | CACCAGTGTTCTCTTGCAGACAT |
| | | Probe-Positive | /56-FAM/AG CCT ACC T/ZEN/T GAC AAG CAA TCA GAC ACT CAA /3IABkFQ/ |
| | | gBlock | GAGGCATCAATATCGGACATGGCTTCGGACAGTCGCTGTCCAACACAAGGTG AAGCCTACCTTGACAAGCAATCAGACACTCAATATGTCTGCAAGAGAACACT GGTGGATAGAGGTTGGGGAAATGGGTGTGGACT |
| Asian lineage | RT-qPCR | qPCR-As-F | CCGCTGCCCAACACAAG |
| | | qPCR-As-R | CCACTAACGTTCTTTTGCAGACAT |
| | | Probe-Positive | Same as above |
| | | gBlock | GAGGCATCAATATCAGACATGGCTTCTGACAGCCGCTGCCCAACACAAGGTGAAGCCT ACCTTGACAAGCAATCAGACACTCAATATGTCTGCAAAAGAACGTTAGTGGACAGAGGC TGGGGAAATGGATGTGGACT |
| **RT-qPCR for *Actin*** | | | |
| *Aedes aegypti* | RT-qPCR | Actin-F | CGTTCGTGACATCAAGGAAA |
| | | Actin-R | GACGGCTGGAAGAGGGC |
| **Strand specific RT-qPCR for ZIKV (genomic sense)** | | | |
| African lineage | RT | Tag-ZIKV-Af-R | GGCCGTCATGGTGGCGAATAACACCAGTGTTCTCTTGCAGACAT |
| | qPCR | ZIKV-Af-F | AATAAATCATAAGTCGCTGTCCAACACAAG |
| | | Tag | AATAAATCATAAGGCCGTCATGGTGGCGAATAA |
| | | Probe-Positive | Same as above |
| Asian lineage | RT | Tag-ZIKV-As-R | GGCCGTCATGGTGGCGAATAACCACTAACGTTCTTTTGCAGACAT |
| | qPCR | ZIKV-As-F | AATAAATCATAACCGCTGCCCAACACAAG |
| | | Tag | Same as above |
| | | Probe-Positive | Same as above |
| **Strand specific RT-qPCR for ZIKV (antigenomic sense)** | | | |
| African lineage | PCR (standard curves) | antiZIKV-Af-st-R | ATCAGGTGCATAGGAGTTAGCAATAGA |
| | | antiZIKV-Af-st-F-T7 | TAA TAC GAC TCA CTA TAG AGCAGAAACAGCCGTGGAAAGG |
| | RT | Tag-ZIKV-Af-F | GGCCGTCATGGTGGCGAATAAGTCGCTGTCCAACACAAG |
| | qPCR | ZIKV-Af-R | AATAAATCATAACACCAGTGTTCTCTTGCAGACAT |
| | | Tag | Same as above |
| | | Probe-Negative | /56-FAM/TT GAG TGT C/ZEN/T GAT TGC TTG TCA AGG TAG GCT /3IABkFQ/ |
| Asian lineage | PCR (standard curves) | antiZIKV- As-st-R | ATCAGGTGCATAGGAGTCAGTAATAGGG |
| | | antiZIKV-As-st-F-T7 | TAA TAC GAC TCA CTA TAG AGCGGAGACGGCTGTGGATAAG |
| | RT | Tag-ZIKV-As-F | GGCCGTCATGGTGGCGAATAACCGCTGCCCAACACAAG |
| | qPCR | ZIKV-As-R | AATAAATCATAACCACTAACGTTCTTTTGCAGACAT |

| | | | |
|---|---|---|---|
| | | Tag | Same as above |
| | | Probe-Negative | Same as above |

Table S3. Raw data of experimental mosquito infections.

| Experiment | *Ae. aegypti* colony | | Prevalence over time | | | | | | | | |
|---|---|---|---|---|---|---|---|---|---|---|---|
| Natural isolates and Reverse generated viruses | Colombia | Infection prevalence | Day 7 | | | Day 14 | | | | | |
| | | Virus | Total | Positive | Ratio | Total | Positive | Ratio | | | |
| | | iSenegal | 18 | 16 | 0.89 | 18 | 17 | 0.94 | | | |
| (Figure S2) | | rSenegal | 19 | 17 | 0.89 | 25 | 22 | 0.88 | | | |
| | | iThailand | 19 | 17 | 0.89 | 27 | 27 | 1.00 | | | |
| | | rThailand | 24 | 22 | 0.92 | 18 | 18 | 1.00 | | | |
| | | Dissemination prevalence | Day 7 | | | Day 14 | | | | | |
| | | Virus | Total | Positive | Ratio | Total | Positive | Ratio | | | |
| | | iSenegal | 16 | 11 | 0.69 | 17 | 11 | 0.65 | | | |
| | | rSenegal | 17 | 7 | 0.41 | 22 | 12 | 0.55 | | | |
| | | iThailand | 17 | 0 | 0.00 | 27 | 5 | 0.19 | | | |
| | | rThailand | 22 | 1 | 0.05 | 18 | 3 | 0.17 | | | |
| | | Transmission prevalence | Day 7 | | | Day 14 | | | | | |
| | | Virus | Total | Positive | Ratio | Total | Positive | Ratio | | | |
| | | Senegal | 11 | 0 | 0.00 | 11 | 7 | 0.64 | | | |
| | | rSenegal | 7 | 0 | 0.00 | 12 | 10 | 0.83 | | | |
| | | Thailand | 0 | 0 | 0.00 | 5 | 0 | 0.00 | | | |
| | | rThailand | 1 | 0 | 0.00 | 3 | 0 | 0.00 | | | |
| 1$^{st}$ panel | Colombia | Infection prevalence | Day 7 | | | Day 10 | | | Day 14 | | |
| (Figure 3A-C) | | Virus | Total | Positive | Ratio | Total | Positive | Ratio | Total | Positive | Ratio |
| | | rSenegal | 34 | 33 | 0.97 | 36 | 35 | 0.97 | 37 | 35 | 0.95 |
| | | rThailand | 30 | 26 | 0.87 | 31 | 28 | 0.90 | 43 | 40 | 0.93 |
| | | Sen/Thai-SP | 45 | 38 | 0.84 | 45 | 44 | 0.98 | 40 | 34 | 0.85 |
| | | Thai/Sen-SP | 44 | 44 | 1.00 | 32 | 32 | 1.00 | 45 | 43 | 0.96 |
| | | Sen/Thai-nSP | 38 | 36 | 0.95 | 46 | 46 | 1.00 | 34 | 34 | 1.00 |
| | | Thai/Sen-nSP | 31 | 29 | 0.94 | 34 | 31 | 0.91 | 26 | 24 | 0.92 |
| | | Sen/Thai-UTR | 39 | 36 | 0.92 | 26 | 24 | 0.92 | 35 | 32 | 0.91 |
| | | Thai/Sen-UTR | 38 | 35 | 0.92 | 32 | 30 | 0.94 | 46 | 45 | 0.98 |
| | | Dissemination prevalence | Day 7 | | | Day 10 | | | Day 14 | | |
| | | Virus | Total | Positive | Ratio | Total | Positive | Ratio | Total | Positive | Ratio |
| | | rSenegal | 33 | 30 | 0.91 | 35 | 35 | 1.00 | 35 | 34 | 0.97 |
| | | rThailand | 26 | 22 | 0.85 | 28 | 28 | 1.00 | 40 | 38 | 0.95 |
| | | Sen/Thai-SP | 38 | 33 | 0.87 | 44 | 38 | 0.86 | 34 | 34 | 1.00 |
| | | Thai/Sen-SP | 44 | 36 | 0.82 | 32 | 31 | 0.97 | 43 | 43 | 1.00 |

| | | | | | | | | | | | |
|---|---|---|---|---|---|---|---|---|---|---|---|
| | | Sen/Thai-nSP | 36 | 34 | 0.94 | 46 | 45 | 0.98 | 34 | 33 | 0.97 |
| | | Thai/Sen-nSP | 29 | 25 | 0.86 | 31 | 30 | 0.97 | 24 | 24 | 1.00 |
| | | Sen/Thai-UTR | 36 | 32 | 0.89 | 24 | 24 | 1.00 | 32 | 32 | 1.00 |
| | | Thai/Sen-UTR | 35 | 31 | 0.89 | 30 | 26 | 0.87 | 45 | 43 | 0.96 |
| | | Transmission prevalence | Day 7 | | | Day 10 | | | Day 14 | | |
| | | Virus | Total | Positive | Ratio | Total | Positive | Ratio | Total | Positive | Ratio |
| | | rSenegal | 30 | 3 | 0.10 | 35 | 18 | 0.51 | 34 | 17 | 0.50 |
| | | rThailand | 22 | 0 | 0.00 | 28 | 1 | 0.04 | 38 | 4 | 0.11 |
| | | Sen/Thai-SP | 33 | 1 | 0.03 | 38 | 4 | 0.11 | 34 | 7 | 0.21 |
| | | Thai/Sen-SP | 36 | 2 | 0.06 | 31 | 9 | 0.29 | 43 | 15 | 0.35 |
| | | Sen/Thai-nSP | 34 | 1 | 0.03 | 45 | 5 | 0.11 | 33 | 8 | 0.24 |
| | | Thai/Sen-nSP | 25 | 0 | 0.00 | 30 | 4 | 0.13 | 24 | 4 | 0.17 |
| | | Sen/Thai-UTR | 32 | 2 | 0.06 | 24 | 7 | 0.29 | 32 | 13 | 0.41 |
| | | Thai/Sen-UTR | 31 | 0 | 0.00 | 26 | 0 | 0.00 | 43 | 2 | 0.05 |
| 2nd panel | Colombia | Infection prevalence | Day 7 | | | Day 10 | | | Day 14 | | |
| (Figure 3D-F) | | Virus | Total | Positive | Ratio | Total | Positive | Ratio | Total | Positive | Ratio |
| | | rSenegal | 26 | 26 | 1.00 | 25 | 25 | 1.00 | 22 | 22 | 1.00 |
| | | Sen/Thai-C | 28 | 28 | 1.00 | 35 | 35 | 1.00 | 32 | 32 | 1.00 |
| | | Sen/Thai-prM | 48 | 47 | 0.98 | 30 | 30 | 1.00 | 15 | 15 | 1.00 |
| | | Sen/Thai-E | 27 | 27 | 1.00 | 28 | 28 | 1.00 | 24 | 24 | 1.00 |
| | | Sen/Thai-NS1 | 26 | 26 | 1.00 | 22 | 22 | 1.00 | 21 | 21 | 1.00 |
| | | Sen/Thai-NS2 | 25 | 24 | 0.96 | 22 | 22 | 1.00 | 31 | 31 | 1.00 |
| | | Sen/Thai-NS3 | 30 | 29 | 0.97 | 22 | 22 | 1.00 | 21 | 21 | 1.00 |
| | | Sen/Thai-NS4 | 38 | 36 | 0.95 | 18 | 18 | 1.00 | 16 | 16 | 1.00 |
| | | Sen/Thai-NS5 | 27 | 27 | 1.00 | 14 | 14 | 1.00 | 10 | 10 | 1.00 |
| | | Dissemination prevalence | Day 7 | | | Day 10 | | | Day 14 | | |
| | | Virus | Total | Positive | Ratio | Total | Positive | Ratio | Total | Positive | Ratio |
| | | rSenegal | 26 | 21 | 0.81 | 25 | 25 | 1.00 | 22 | 22 | 1.00 |
| | | Sen/Thai-C | 28 | 27 | 0.96 | 35 | 35 | 1.00 | 32 | 32 | 1.00 |
| | | Sen/Thai-prM | 47 | 42 | 0.89 | 30 | 30 | 1.00 | 15 | 15 | 1.00 |
| | | Sen/Thai-E | 27 | 24 | 0.89 | 28 | 28 | 1.00 | 24 | 24 | 1.00 |
| | | Sen/Thai-NS1 | 26 | 25 | 0.96 | 22 | 22 | 1.00 | 21 | 21 | 1.00 |
| | | Sen/Thai-NS2 | 24 | 24 | 1.00 | 22 | 22 | 1.00 | 31 | 31 | 1.00 |
| | | Sen/Thai-NS3 | 29 | 29 | 1.00 | 22 | 22 | 1.00 | 21 | 21 | 1.00 |
| | | Sen/Thai-NS4 | 36 | 34 | 0.94 | 18 | 18 | 1.00 | 16 | 16 | 1.00 |
| | | Sen/Thai-NS5 | 27 | 27 | 1.00 | 14 | 14 | 1.00 | 10 | 9 | 0.90 |
| | | Transmission prevalence | Day 7 | | | Day 10 | | | Day 14 | | |

| | | | | | | | | | | | |
|---|---|---|---|---|---|---|---|---|---|---|---|
| | | Virus | Total | Positive | Ratio | Total | Positive | Ratio | Total | Positive | Ratio |
| | | rSenegal | 21 | 3 | 0.14 | 25 | 4 | 0.16 | 22 | 12 | 0.55 |
| | | Sen/Thai-C | 27 | 1 | 0.04 | 35 | 7 | 0.20 | 32 | 17 | 0.53 |
| | | Sen/Thai-prM | 42 | 7 | 0.17 | 30 | 0 | 0.00 | 15 | 5 | 0.33 |
| | | Sen/Thai-E | 24 | 1 | 0.04 | 28 | 2 | 0.07 | 24 | 7 | 0.29 |
| | | Sen/Thai-NS1 | 25 | 4 | 0.16 | 22 | 2 | 0.09 | 21 | 8 | 0.38 |
| | | Sen/Thai-NS2 | 24 | 1 | 0.04 | 22 | 3 | 0.14 | 31 | 22 | 0.71 |
| | | Sen/Thai-NS3 | 29 | 3 | 0.10 | 22 | 5 | 0.23 | 21 | 15 | 0.71 |
| | | Sen/Thai-NS4 | 34 | 3 | 0.09 | 18 | 2 | 0.11 | 16 | 5 | 0.31 |
| | | Sen/Thai-NS5 | 27 | 2 | 0.07 | 14 | 4 | 0.29 | 9 | 4 | 0.44 |
| 3rd panel | Colombia | Infection prevalence | Day 7 | | | Day 10 | | | Day 14 | | |
| (Figure 3G-I) | | Virus | Total | Positive | Ratio | Total | Positive | Ratio | Total | Positive | Ratio |
| | | rThailand | 18 | 17 | 0.94 | 20 | 19 | 0.95 | 17 | 16 | 0.94 |
| | | Thai/Sen-C | 26 | 25 | 0.96 | 28 | 27 | 0.96 | 20 | 20 | 1.00 |
| | | Thai/Sen-prM | 16 | 15 | 0.94 | 26 | 22 | 0.85 | 20 | 17 | 0.85 |
| | | Thai/Sen-E | 20 | 19 | 0.95 | 25 | 25 | 1.00 | 23 | 23 | 1.00 |
| | | Thai/Sen-NS1 | 18 | 17 | 0.94 | 25 | 24 | 0.96 | 19 | 18 | 0.95 |
| | | Thai/Sen-NS2 | 22 | 18 | 0.82 | 25 | 24 | 0.96 | 27 | 23 | 0.85 |
| | | Thai/Sen-NS3 | 33 | 26 | 0.79 | 23 | 21 | 0.91 | 32 | 26 | 0.81 |
| | | Thai/Sen-NS4 | 18 | 16 | 0.89 | 22 | 16 | 0.73 | 35 | 27 | 0.77 |
| | | Thai/Sen-NS5 | 25 | 23 | 0.92 | 26 | 20 | 0.77 | 23 | 18 | 0.78 |
| | | Dissemination prevalence | Day 7 | | | Day 10 | | | Day 14 | | |
| | | Virus | Total | Positive | Ratio | Total | Positive | Ratio | Total | Positive | Ratio |
| | | rThailand | 17 | 15 | 0.88 | 19 | 18 | 0.95 | 16 | 14 | 0.88 |
| | | Thai/Sen-C | 25 | 24 | 0.96 | 27 | 22 | 0.81 | 20 | 19 | 0.95 |
| | | Thai/Sen-prM | 15 | 12 | 0.80 | 22 | 17 | 0.77 | 17 | 14 | 0.82 |
| | | Thai/Sen-E | 19 | 18 | 0.95 | 25 | 24 | 0.96 | 23 | 22 | 0.96 |
| | | Thai/Sen-NS1 | 17 | 10 | 0.59 | 24 | 23 | 0.96 | 18 | 18 | 1.00 |
| | | Thai/Sen-NS2 | 18 | 14 | 0.78 | 24 | 22 | 0.92 | 23 | 23 | 1.00 |
| | | Thai/Sen-NS3 | 26 | 17 | 0.65 | 21 | 21 | 1.00 | 26 | 24 | 0.92 |
| | | Thai/Sen-NS4 | 16 | 5 | 0.31 | 16 | 16 | 1.00 | 27 | 27 | 1.00 |
| | | Thai/Sen-NS5 | 23 | 21 | 0.91 | 20 | 18 | 0.90 | 18 | 18 | 1.00 |
| | | Transmission prevalence | Day 7 | | | Day 10 | | | Day 14 | | |
| | | Virus | Total | Positive | Ratio | Total | Positive | Ratio | Total | Positive | Ratio |
| | | rThailand | 15 | 0 | 0 | 18 | 0 | 0 | 14 | 0 | 0.00 |
| | | Thai/Sen-C | 24 | 0 | 0 | 22 | 0 | 0 | 19 | 4 | 0.21 |
| | | Thai/Sen-prM | 12 | 0 | 0 | 17 | 0 | 0 | 14 | 0 | 0.00 |

| | | | | | | | | | | | |
|---|---|---|---|---|---|---|---|---|---|---|---|
| | | Thai/Sen-E | 18 | 0 | 0 | 24 | 0 | 0 | 22 | 4 | 0.18 |
| | | Thai/Sen-NS1 | 10 | 0 | 0 | 23 | 0 | 0 | 18 | 2 | 0.11 |
| | | Thai/Sen-NS2 | 14 | 0 | 0 | 22 | 0 | 0 | 23 | 1 | 0.04 |
| | | Thai/Sen-NS3 | 17 | 0 | 0 | 21 | 0 | 0 | 24 | 2 | 0.08 |
| | | Thai/Sen-NS4 | 5 | 0 | 0 | 16 | 0 | 0 | 27 | 2 | 0.07 |
| | | Thai/Sen-NS5 | 21 | 0 | 0 | 18 | 0 | 0 | 18 | 3 | 0.17 |
| 1st panel | Cape Verde | Probability of infection | | Day 3 | | | | | | | |
| (Figure S3) | | Virus | Dose (PFU/ml) | Total | Positive | Ratio | | | | | |
| | | rSenegal | 1300000 | 21 | 20 | 0.95 | | | | | |
| | | | 50000 | 37 | 32 | 0.86 | | | | | |
| | | | 14000 | 29 | 14 | 0.48 | | | | | |
| | | rThailand | 700000 | 27 | 20 | 0.74 | | | | | |
| | | | 40000 | 28 | 4 | 0.14 | | | | | |
| | | | 20000 | 27 | 0 | 0.00 | | | | | |
| | | Sen/Thai-SP | 1000000 | 25 | 23 | 0.92 | | | | | |
| | | | 60000 | 27 | 9 | 0.33 | | | | | |
| | | | 10000 | 42 | 1 | 0.02 | | | | | |
| | | Thai/Sen-SP | 1050000 | 26 | 20 | 0.77 | | | | | |
| | | | 40000 | 26 | 11 | 0.42 | | | | | |
| | | | 10000 | 29 | 2 | 0.07 | | | | | |
| | | Sen/Thai-nSP | 6000000 | 19 | 19 | 1.00 | | | | | |
| | | | 2000000 | 7 | 6 | 0.86 | | | | | |
| | | | 200000 | 20 | 7 | 0.35 | | | | | |
| | | Thai/Sen-nSP | 1600000 | 25 | 21 | 0.84 | | | | | |
| | | | 100000 | 36 | 5 | 0.14 | | | | | |
| | | | 20000 | 31 | 0 | 0.00 | | | | | |
| | | Sen/Thai-UTR | 1050000 | 26 | 26 | 1.00 | | | | | |
| | | | 80000 | 27 | 19 | 0.70 | | | | | |
| | | | 6000 | 27 | 7 | 0.26 | | | | | |
| | | Thai/Sen-UTR | 2000000 | 10 | 7 | 0.70 | | | | | |
| | | | 60000 | 25 | 1 | 0.04 | | | | | |
| | | | 10000 | 29 | 0 | 0.00 | | | | | |

### Table S4. Statistical analysis of viral infection, dissemination and transmission in mosquitoes.

The table shows the results of the multivariate logistic regression of the proportion of blood-fed mosquitoes with a midgut infection (infection prevalence), the proportion of midgut-infected mosquitoes with a disseminated infection (dissemination prevalence), and the proportion of mosquitoes with a disseminated infection that expectorated virus in their saliva (transmission prevalence). The colony of mosquitoes and the panel of chimeric viruses are indicated. For each phenotype, the minimal adequate model was obtained by sequentially removing non-significant terms ($p$ <0.05) from the full-factorial model. Df: degrees of freedom; LR: likelihood ratio.

| | | | | Infection | | Dissemination | | Transmission | |
|---|---|---|---|---|---|---|---|---|---|
| *Ae. aegypti* colony | Chimeric viruses | Factor | Df | LR $Chi^2$ | *P* value | LR $Chi^2$ | *P* value | LR $Chi^2$ | *P* value |
| Colombia | 1st panel (Figure 3A-C) | Experiment | 1 | | | 18.43 | <0.0001 | 6.006 | 0.0143 |
| | | Virus | 7 | 34.98 | <0.0001 | 17.34 | 0.0153 | 69.35 | <0.0001 |
| | | Time | 1 | | | | | 26.13 | <0.0001 |
| | 2nd panel (Figure 3D-F) | Experiment | 1 | | | 18.53 | <0.0001 | 11.59 | 0.0007 |
| | | Virus | 8 | | | <0.0001 | 1 | 16.87 | 0.0315 |
| | | Time | 1 | 9.152 | 0.0025 | 0.0004 | 0.9844 | 102.3 | <0.0001 |
| | | Experiment*Time | 1 | | | | | 4.102 | 0.0428 |
| | | Virus*Time | 8 | | | 17.75 | 0.0232 | | |
| | 3rd panel (Figure 3G-I) | Virus | 8 | 34.51 | <0.0001 | 16.89 | 0.0313 | | |
| | | Time | 1 | | | 31.86 | <0.0001 | 37.9 | <0.0001 |
| | | Virus*Time | 8 | | | 36.82 | <0.0001 | | |

Table S5. Raw data of experimental mosquito infections and statistical analysis of transmission efficiency in mosquitoes.

The table shows the results of the multivariate logistic regression of the proportion of blood-fed mosquitoes that expectorated virus in their saliva (transmission efficiency). The colony of mosquitoes and the panel of chimeric viruses are indicated. The minimal adequate model was obtained by sequentially removing non-significant terms (*p* <0.05) from the full-factorial model. Df: degrees of freedom; LR: likelihood ratio.

| Experiment | *Ae. aegypti* colony | Transmission efficiency | | | | | | | | | | Statistical analysis | | | |
|---|---|---|---|---|---|---|---|---|---|---|---|---|---|---|---|
| 1st panel (Figure S5A) | Uganda | | Day 7 | | | Day 10 | | | Day 14 | | | Factor | Df | LR $Chi^2$ | *P* value |
| | | Virus | Total | Positive | Ratio | Total | Positive | Ratio | Total | Positive | Ratio | Virus | 7 | 34.65 | <0.0001 |
| | | rSenegal | 11 | 2 | 0.18 | 20 | 5 | 0.25 | 20 | 6 | 0.30 | Time | 1 | <0.0001 | 0.9980 |
| | | rThailand | 16 | 0 | 0.00 | 17 | 0 | 0.00 | 16 | 0 | 0.00 | | | | |
| | | Sen/Thai-SP | 14 | 0 | 0.00 | 18 | 1 | 0.06 | 22 | 2 | 0.09 | | | | |
| | | Thai/Sen-SP | 10 | 0 | 0.00 | 14 | 0 | 0.00 | 22 | 5 | 0.23 | | | | |
| | | Sen/Thai-nSP | 14 | 0 | 0.00 | 17 | 2 | 0.12 | 18 | 3 | 0.17 | | | | |
| | | Thai/Sen-nSP | 27 | 0 | 0.00 | 27 | 1 | 0.04 | 28 | 7 | 0.25 | | | | |
| | | Sen/Thai-UTR | 12 | 0 | 0.00 | 21 | 0 | 0.00 | 29 | 12 | 0.41 | | | | |
| | | Thai/Sen-UTR | 15 | 0 | 0.00 | 18 | 0 | 0.00 | 22 | 0 | 0.00 | | | | |
| 1st panel (Figure S5B) | Cape Verde | | Day 7 | | | Day 10 | | | Day 14 | | | Factor | Df | LR $Chi^2$ | *P* value |
| | | Virus | Total | Positive | Ratio | Total | Positive | Ratio | Total | Positive | Ratio | Virus* Time | 7 | 14.81 | 0.0384 |
| | | rSenegal | 27 | 2 | 0.07 | 24 | 9 | 0.38 | 25 | 13 | 0.52 | Virus | 7 | 89.40 | <0.0001 |
| | | rThailand | 22 | 0 | 0.00 | 26 | 0 | 0.00 | 27 | 1 | 0.04 | Time | 1 | 51.58 | <0.0001 |
| | | Sen/Thai-SP | 22 | 0 | 0.00 | 25 | 3 | 0.12 | 18 | 5 | 0.28 | | | | |
| | | Thai/Sen-SP | 17 | 0 | 0.00 | 21 | 1 | 0.05 | 28 | 8 | 0.29 | | | | |
| | | Sen/Thai-nSP | 20 | 0 | 0.00 | 20 | 0 | 0.00 | 19 | 5 | 0.26 | | | | |
| | | Thai/Sen-nSP | 28 | 0 | 0.00 | 24 | 1 | 0.04 | 29 | 1 | 0.03 | | | | |
| | | Sen/Thai-UTR | 21 | 1 | 0.05 | 25 | 9 | 0.36 | 22 | 12 | 0.55 | | | | |
| | | Thai/Sen-UTR | 29 | 0 | 0.00 | 24 | 0 | 0.00 | 20 | 0 | 0.00 | | | | |
